# AI-driven discovery and engineering of human endogenous nanocage proteins for mRNA delivery

**DOI:** 10.64898/2026.02.07.704552

**Authors:** Dongmin Xun, Lanqing Li, Yi Liu, Ziyue Yang, Jie Zou, Qianyi Liang, Ke Li, Zhu Zhu, Jicong Zhang, Lijun Lang, Kun Liu, Xinyu Wang, Meilin Tian, Guangyong Chen, Bolin An, Pheng Ann Heng, Chao Zhong

## Abstract

Safe and effective gene delivery remains a central challenge for therapeutic applications. While non-viral and viral vectors have enabled substantial progress, their reliance on non-human components often triggers immune responses, limiting their use in chronic treatments. Here, we developed DeepDelivery, an artificial intelligence-driven platform to repurpose human proteins for mRNA delivery. An unbiased screening of the human proteome nominated 512 candidates, with experimental validation confirming that 80% of top-ranked hits form mRNA-encapsulating particles and mediate efficient functional delivery in human cells without provoking detectable inflammation. Notably, multiple tripartite motif (TRIM) family proteins, typically linked to antiviral responses, exhibited strong assembly and delivery activity. Quantitative analysis and interpretation of the model revealed structural domains that govern nanocage formation, enabling domain-guided engineering of TRIM25 variants with enhanced function. Our work establishes a generalizable framework for discovering human-derived delivery vehicles and provides a path toward programmable, non-immunogenic mRNA therapeutics.

## Main text

Protein-based delivery systems represent a promising class of therapeutic carriers due to their biocompatibility, structural programmability, and modular assembly properties ^1-3^. Among these, virus-like particles (VLPs) have been widely adopted for nucleic acid and protein encapsulation ^4,5^. These architectures are typically derived from viral capsid proteins that have been stripped of replicative elements and engineered to package therapeutic cargo ^6-11^. Recent advances demonstrate their utility in delivering complex molecular payloads, including CRISPR-Cas systems and mRNA, across diverse biological settings ^12-16^. Similarly, synthetic and enveloped protein nanocages have shown efficacy in mediating intercellular RNA transfer ^17^, highlighting the broader potential of self-assembling protein systems in biomedicine.

Despite this progress, most current delivery platforms rely on non-human proteins, which pose risks of immunogenicity and may limit clinical translation ^18^. Repeated administration of viral components can provoke adaptive immune responses, further restricting dosing regimens and therapeutic utility ^19^. These challenges have spurred interest in identifying human proteins capable of self-assembly as safer and more biocompatible alternatives. A handful of endogenous human proteins, such as Arc, a retrotransposon-derived neuronal protein that packages RNA into extracellular capsids ^20,21^, and PEG10, which naturally forms mRNA-encapsulating particles and has been engineered for RNA delivery ^4^, illustrate the existence of native nanocage-forming scaffolds. Yet, systematic discovery remains constrained by methodological limitations. Conventional homology-based searches are biased toward retroelement-derived genes and often miss proteins with divergent or cryptic assembly domains ^22,23^. For example, a genome-wide survey has identified only a limited number of capsid-like human proteins, with PEG10 representing a rare functional example ^24^. While de novo protein design can generate novel assemblies ^25^, it is less suited for detecting innate self-assembling proteins with complex sequence-structure relationships. Consequently, the full scope of human proteins capable of forming nanocages remains largely unexplored.

To address this gap, we developed DeepDelivery, an AI-driven pipeline that combines large-scale predictive modeling with experimental validation (Fig.1a). Neural networks generated by the DeepDelivery model factory were trained on self-curated proprietary datasets to learn sequence and structural features predictive of self-assembly. These models were then applied to the entire human-expressible proteome using a dual-model screening strategy designed to maximize both sensitivity and specificity, ultimately identifying 512 proteins with predicted nanocage-forming potential. Experimental testing of 15 top-ranked candidates showed that 12 formed mRNA-encapsulating particles in cells. A UMAP (Uniform Manifold Approximation and Projection) visualization of the human-expressible proteome revealed that DeepDelivery-predicted positives cluster in a small area of the embedding space together with the previously reported VLP-forming proteins Arc and PEG10, indicating that the model captures shared features linking newly identified candidates to known VLP-forming proteins (Fig. 1b-f). This approach extends beyond conventional motif-based detection by capturing higher-order sequence patterns associated with protein assembly ^19,26-29^. Notably, several of the predicted candidates belong to the TRIM family, which is classically associated with innate immunity. Interpretable AI analyses indicated that the conserved RBCC (RING, B-box, coiled-coil) domains promote higher-order assembly. Guided by these insights, we engineered TRIM variants with improved mRNA-delivery efficiency, illustrating the potential for rational optimization of human-derived nanocages. Together, our findings establish a systematic, generalizable strategy for discovering and engineering endogenous human proteins with self-assembly capacity, paving the way for a new class of immunocompatible and customizable delivery systems for therapeutic applications.

**Fig. 1.**
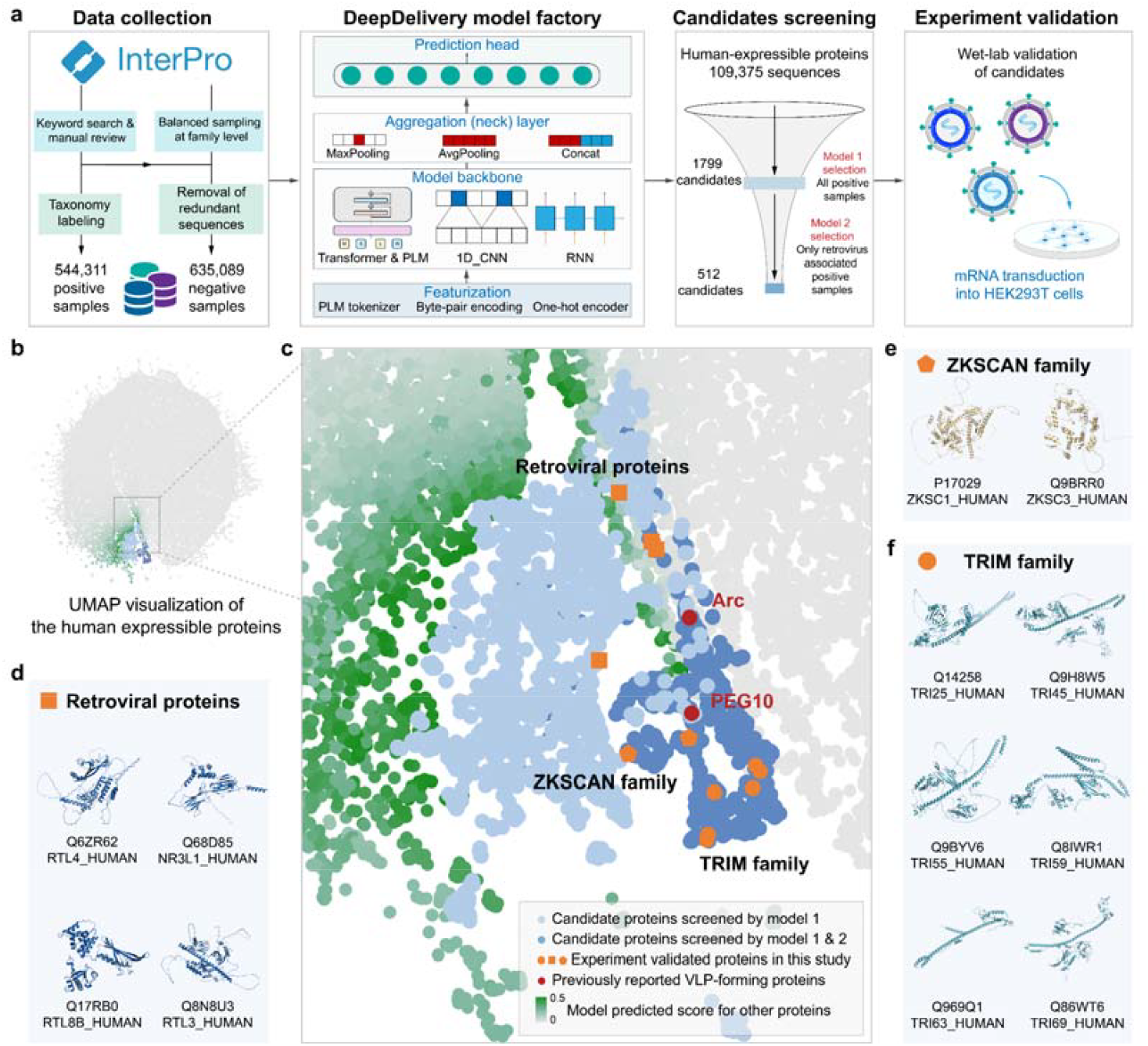
AI-assisted discovery of human protein nanocages. **a**, DeepDelivery workflow, where AI models generated by the DeepDelivery model factory are trained based on a self-curated balanced dataset consisting of over 1 million nanocage-positive and negative samples in total, along with a subset containing only retroviral capsid proteins and balanced negatives. The deep learning architecture employs a stack of featurization (e.g., Protein Language Model (PLM) tokenizer, byte-pair encoding, or one-hot encoder), model backbone (e.g., PLM, transformer, CNN, or RNN), aggregation (e.g., MaxPool, AvgPool or Concat) layers for systematic implementation of a broad spectrum of AI models, followed by a classification head that assigns each protein a probability score for nanocage-forming potential. After training and selecting the best-performing model architecture, a dual-model screening process was applied to the human proteome (UniProt) of 109,375 sequences, where Model 1 (trained on all positive and negative samples) predicted 1,799 candidates, which were further narrowed down to 512 high-confidence candidates by Model 2 (trained on retroviral capsid proteins and balanced negatives). The top-ranked candidates were chosen for experimental validation of self-assembly and mRNA-delivery capability. **b**, UMAP projection of the human proteome highlighting model-selected candidates (colour legend in c). **c**, Zoomed-in UMAP view showing experimentally validated proteins that form mRNA-transducing nanocages. Red, previously reported cage-forming proteins; light blue, candidates selected by Model 1; blue, candidates selected by both models; orange, experimentally validated proteins from retroviral, ZKSCAN, and TRIM families. **d–f**, Predicted structures of representative candidate proteins using AlphaFold (default parameters; accessed via https://golgi.sandbox.google.com/). **d**, Retroviral proteins. **e**, ZKSCAN family proteins. **f**, TRIM family proteins.

### Curation of Large-Scale Datasets with Nanocage Assembly Annotations

Inspired by pre-existing mRNA nanocage designs, which often exploit the self-assembly properties of viral capsid proteins ^4,17,30^, especially retrovirus capsid proteins, we constructed comprehensive sequence datasets of viral capsid proteins and retrovirus-related proteins to enable robust, data-driven discovery of human nanocage proteins.

Our full dataset (V1) was developed using a curation strategy based on the InterPro database, which consolidates functional annotations from Pfam, TIGRFAMs, and CATH-Gene3D (Supplementary Fig.1a). We first conducted systematic keyword searches (e.g., “capsid,” “viral coat”) to identify protein families potentially involved in VLP formation, resulting in 472 initial candidates. These candidates were then manually reviewed to retain families explicitly annotated as capsid or self-assembling proteins (e.g., IPR000574: Tymovirus coat protein; IPR003309: SCAN domain similar to HIV capsid domain), while excluding unrelated entries (e.g., PF11602: Terminase large subunit). This yielded 456 high-confidence families spanning 68 viral families within the ICTV taxonomy, from *Adenoviridae* to *Virgaviridae*, ultimately encompassing 544,311 non-redundant protein sequences (Fig.1a, Supplementary Fig.2 and 3). We employed the V1 dataset to train our Model 1 for the initial coarse-grained screening. To further refine the candidate pool and to enhance sensitivity toward retrovirus-like motifs, including those present in endogenous retroelements, we created our V2 dataset as a subset of V1 with only retrovirus-derived sequences as positives. The V2 dataset was subsequently used to train our Model 2 to perform a second round of fine-grained filtering (Fig.1a). In the early stage of development, we also explored datasets comprised exclusively of transposon-derived DNA or natural capsid protein sequences, yet to find limited generalization of the trained model on unseen data (Supplementary Fig.1b-c). More discussion of these initial attempts is included in the Supplementary Method section.

Short protein sequences (<100 amino acids) often yield unreliable model predictions due to limited structural context ^31,32^, whereas longer sequences (>1024 amino acids) often exceed input limits of modern protein language models ^33-35^ and may carry redundant information, increasing computational cost with marginal gains in performance. To balance model compatibility, biological relevance, and computational efficiency, we retained sequences longer than 100 residues and truncated those exceeding 1024 amino acids. Sequences shorter than the maximum length were padded to 1024 tokens. Concurrently, we curated an extensive negative control dataset to mitigate potential false positives. Over 18,000 InterPro entries unrelated to capsid formation or self-assembly were selected, encompassing an initial pool of approximately 68 million candidate sequences, which was then randomly sampled, followed by sequence clustering via Diamond BLASTP (e-value < 1×10^-5^, identity > 80%), removing near-duplicate sequences and resulted in 635,089 negative samples. The final V1 dataset achieved an approximate balance between positive and negative classes, providing a high-quality, biologically diverse resource for training and benchmarking deep learning models.

### Deep Learning Model Design and Performance Evaluation

To perform model training and selection, we partitioned the V1 dataset into training, validation, and test sets using two strategies: random split and a cluster-wise split based on 30% sequence identity. These datasets were used to train a variety of deep learning models generated by the DeepDelivery model factory, including 1D-CNN, CNN–RNN, Transformer, and pretrained protein language models (ESM2_t33_650M and ESM2_t6_8M). All models were benchmarked using multiple performance metrics, and the best-performing architecture was selected for subsequent proteome-wide candidate screening (Fig.2a).

**Fig. 2.**
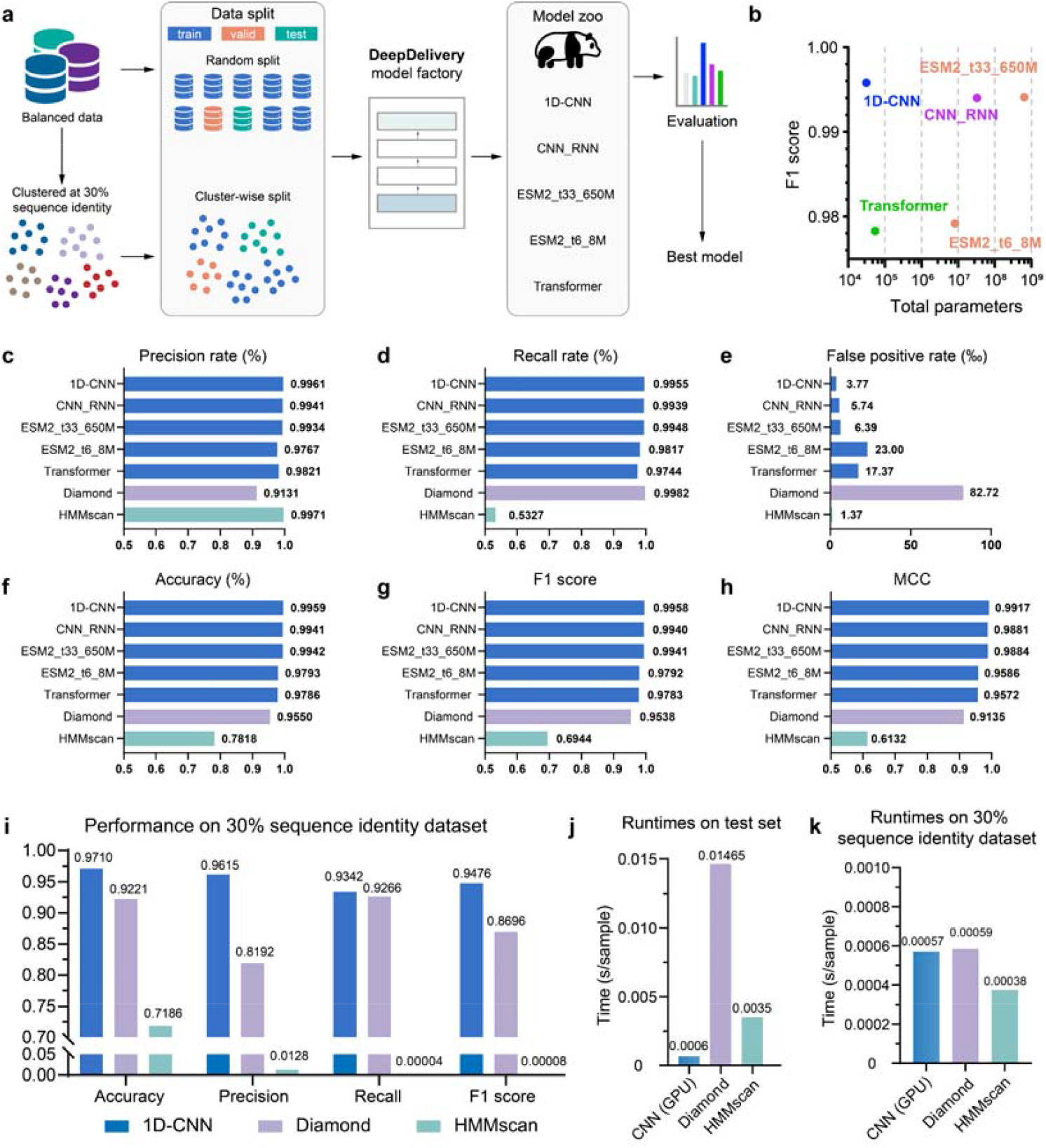
Performance benchmarking of deep learning models and conventional bioinformatics tools for predicting cage-forming proteins. **a**, Positive and negative sequences in the datasets were balanced and partitioned into training, validation, and test sets using two complementary strategies: a standard random split and a cluster-wise split after clustering sequences at 30% sequence identity. Both types of splits were used to train a variety of models (“model zoo”) generated by the DeepDelivery model factory, comprising 1D-CNN, CNN–RNN, Transformer, and pretrained protein language models (ESM2_t33_650M and ESM2_t6_8M). All models were benchmarked on the validation and test sets using performance metrics in **c-k**, and the best-performing architecture was selected for subsequent proteome-wide candidate screening. **b**, Model performance (F1-score) versus complexity (number of parameters, logarithmic scale on the xlaxis). The 1D-CNN model achieves near-optimal prediction performance with the fewest parameters among leading models. **c-h**, Comparative performance across six evaluation metrics: **c**, precision; **d**, recall; **e**, false positive rate; **f**, accuracy; **g**, F1 score; and **h**, Matthew’s correlation coefficient (MCC). **i**, Predictive performance of the 1D-CNN model, Diamond, and HMMscan on a test set produced by the cluster-wise split with 30% sequence identity cutoff. **j-k**, Computational runtime comparison of 1D-CNN, Diamond, and HMMscan on **j** the standard test set and **k** the 30% identity test set.

Model performance was evaluated using a comprehensive suite of metrics: precision, recall, false positive rate (FP rate), accuracy, F1 score, Matthew’s correlation coefficient (MCC), and confusion matrices. To account for potential class imbalance, model selection was based on balanced accuracy and balanced F1 score, in alignment with the ImDrug benchmark guideline^38^. Additional continuous metrics such as area under the receiver operating characteristic curve (ROC-AUC) and area under the precision-recall curve (PR-AUC) were also reported (Supplementary Fig.4-6).

For efficient high-throughput screening, we opted for an optimal balance between predictive performance and computational efficiency while performing model selection. Although larger models (e.g., ESM-2) achieved competitive results in certain metrics, their parameter counts (e.g., ∼2000× that of 1D-CNN) and inference costs rendered them less ideal for computation-sensitive applications. By comparing F1 score against model size on a logarithmic scale (Fig.2b), we identified the 1D-CNN as the ideal option, delivering near-peak performance with minimal complexity. Visualizations via t-SNE further confirmed that 1D-CNN effectively separated positive and negative samples in a high-dimensional space (Supplementary Fig.7). The architectural simplicity of 1D-CNN also facilitates interpretability through methods such as layer-wise relevance propagation (LRP)^39^, enabling identification of protein domains pivotal in predictions and supporting subsequent modular engineering of nanocage candidates.

Previous approaches have largely relied on classical computational techniques^4,24^, such as sequence alignment, to search for homologs of known capsid proteins. To benchmark our deep learning models, we evaluated two widely used bioinformatic tools—Diamond^22^ and HMMscan^23^—using the same V1 dataset and evaluation metrics. All deep learning models consistently outperformed conventional methods across major evaluation metrics (Fig.2c-h), attaining recall and precision rates above 97.7% and 97.4%, respectively. Diamond showed high recall (99.8%) but lower precision (91.3%), resulting in significantly higher false positive rates (<2.3% vs. 8.3%). HMMscan performed poorly, with a recall as low as 53.3%, and correspondingly low F1 and MCC values (Fig.2c-h).

Traditional tools are known to struggle with low-similarity sequences^22,40,41^. To assess model robustness, we clustered the V1 dataset at 30% sequence identity and performed a strict cluster-wise split into training, validation, and test sets. Under this regime, Diamond’s recall dropped from 99.8% to 92.7% (Fig.2d and i) and precision declined from 91.3% to 81.9% (Fig.2c and i), reducing its F1 score by 8.4% (from 95.4% to 87.0%) (Fig.2g and 2i). HMMscan’s performance nearly collapsed, with recall and precision below 1% (Fig.2i). In contrast, a re-trained 1D-CNN model maintained robust performance across all metrics even under data distribution shifts (recall: 93.4%, precision: 96.2%, F1: 94.8%) (Fig.2i).

For assessment of computational efficiency, runtime analysis (Fig.2j) showed that when processing the V3 dataset on a NVIDIA V100 GPU, 1D-CNN was approximately 25 times faster than Diamond v2.1.11 and 6 times faster than HMMscan v.3.2.2. Considering that the runtime performance of Diamond and HMMscan varied depending on dataset composition, further runtime analysis on low similarity datasets (Fig.2k and Supplementary Fig.8) confirmed that 1D-CNN achieved superior predictive accuracy with consistently higher computational efficiency compared to classical bioinformatics methods.

### Screening the Human Proteome for Cage-Forming Protein Candidates

We next applied the selected 1D-CNN models to screen the human proteome for endogenous proteins capable of forming nanocage structures. The UniProt dataset was filtered using the organism label “Homo sapiens (Human)”, yielding 109,375 protein entries with lengths between 100 and 1024 amino acids (98,043 protein entries after removing duplicates) as of October 12, 2023. To improve prediction robustness, we implemented a dual-model screening strategy. Model 1 was trained on a comprehensive dataset encompassing diverse viral families to capture general features associated with capsid formation. Model 2 was trained exclusively on retrovirus-derived sequences, with a balanced set of negatives (∼1:1 ratio), to enhance sensitivity toward retrovirus-like motifs, including those present in endogenous retroelements. A protein was designated as a nanocage-forming candidate only if both models simultaneously classified it as positive, thereby increasing confidence and reducing the false positive rate.

Each candidate sequence received a nanocage-formation probability score. Using a confidence threshold of 0.5 and maintaining the 100-1024 amino acid length constraint, model 1 generated an initial pool of 1799 proteins, in which 512 candidates were further identified by model 2, accounting for 0.47% of the screened proteome (Fig.3a). Among these, several previously documented human cage-forming proteins, including Arc^20,21^, PEG10^4^, were successfully recalled. We further annotated the expression profiles of these candidates using tissue- and cell-type-specific data from the Human Protein Atlas (https://www.proteinatlas.org/). Immune cells showed the highest number of expressed candidates (107), followed by germ cells (67), neurons (62), cancer cells (18), hepatocytes (16), and other cell types. An additional 112 candidates exhibited ubiquitous expression across multiple tissues (Fig.3b and Supplementary Fig.9).

**Fig. 3.**
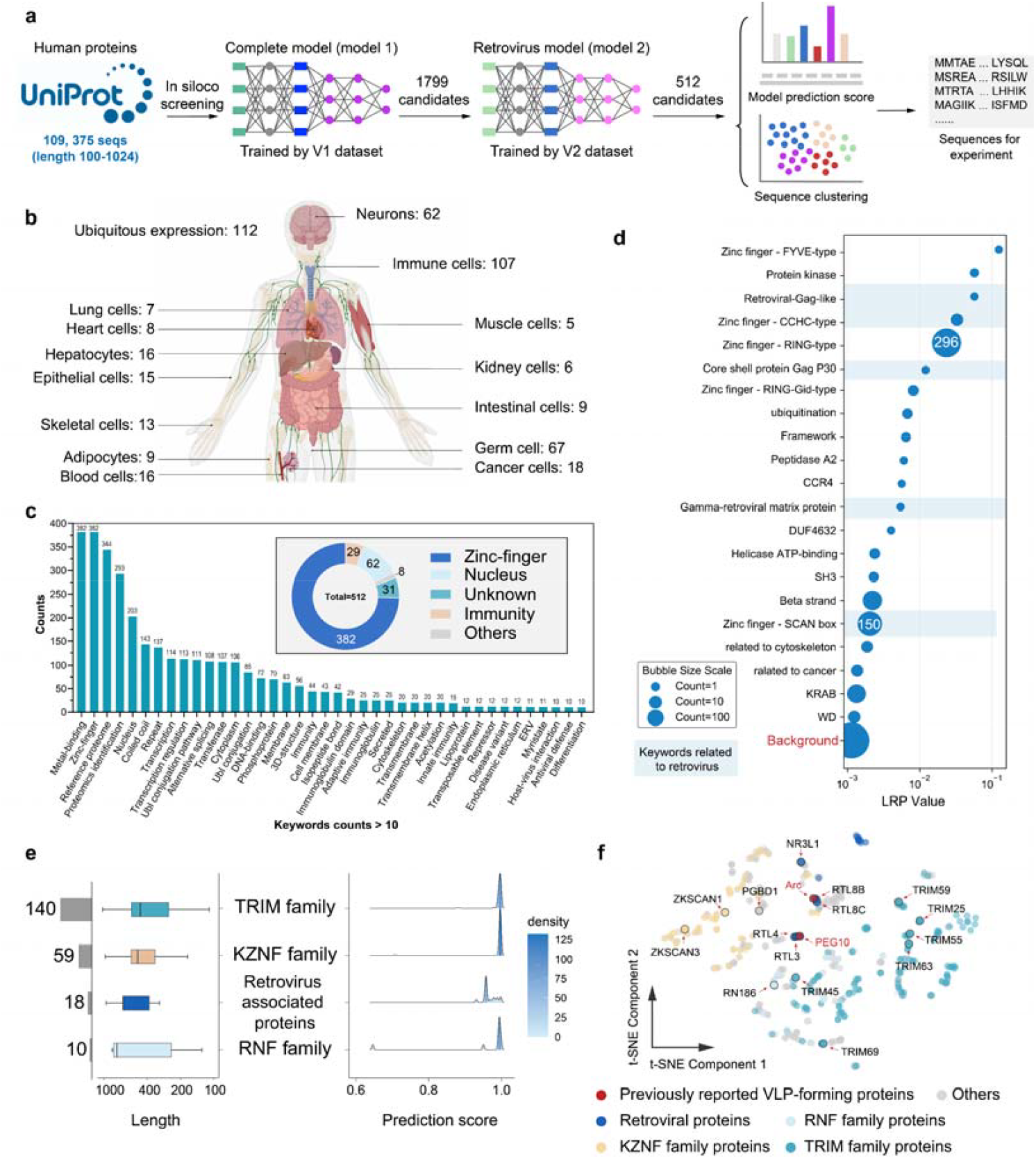
Computational screening and characterization of candidate nanocage-forming proteins from the human proteome. **a**, Screening workflow. Human proteins from UniProtKB were analyzed using two 1D-CNN-based models: a general capsid prediction model and a retrovirus-specific model. 512 high-confidence candidates were identified and prioritized through model scoring and sequence clustering. **b**, Tissue expression profile of the 512 candidates, annotated using data from the Human Protein Atlas. **c**, Functional enrichment analysis based on UniProt keyword annotations (top-right: major functional categories). **d**, LRP analysis of protein domains among all candidates. Retroviral-associated domains are highlighted in blue. Bubble size reflects domain frequency. **e**, Prediction scores and sequence lengths for the top five enriched protein families. Left: score distribution per family. Right: sequence length distribution (light blue bars indicate protein count). **f**, t-SNE projection of model-derived embeddings for all candidates, colored by protein family: known cage-forming (red), TRIM (cyan), KZNF (yellow), RNF (light blue), retroviral (blue), and other (gray). Black circles indicate candidates selected for experimental validation.

Functional annotation of the candidate set using UniProt keywords revealed significant enrichment in terms associated with viral processes, immune response, and nucleic acid binding. Among the 512 candidates, 382 belonged to zinc-finger proteins, 62 were nucleus-associated, 29 were immunity-related, 8 had other functional annotations, and 31 were uncharacterized. Comparative keyword frequency analysis between the candidate set and the full UniProt database showed pronounced enrichment of terms including “zinc-finger,” “adaptive immunity,” “immunity,” and “ERV (endogenous retrovirus)” (Fig.3c and Supplementary Fig.10). Notably, the term “ERV” appeared over 7800 times more frequently among candidates than in the background proteome, suggesting a strong link between retroviral heritage and nanocage-forming potential.

To quantitatively illustrate how protein domains affect model predictions, we applied Layer-wise Relevance Propagation (LRP^39,42^) to identify residues and domains critical for the classification outcome (Fig.3d). Domains with high relevance scores included known capsid-associated motifs such as “Retroviral-Gag-like”, “Core shell protein Gag P30”, and “Gamma-retroviral matrix protein”. Also prominent were nucleic acid-binding domains such as “Zinc finger-CCHC” and “Zinc finger-SCAN box”, the latter sharing structural homology with the C-terminal domain of the HIV capsid protein^43^. These findings corroborate the functional relevance of the candidate set to nanocage assembly.

We further categorized the 512 candidates based on protein family annotations (Fig.3e). The tripartite motif (TRIM) family was most represented (140 members), followed by Kruppel-associated box domain zinc finger proteins (KZNF, 59), retroviral proteins (18), and ring finger proteins (RNF, 10). From these, we selected 17 high-scoring candidates for experimental validation (Supplementary Table 1). Fourteen were drawn from the most abundant families (TRIM, KZNF, RNF, and retroviral proteins), one represented transposable element-derived proteins (PGBD1), and two were previously established nanocage-forming proteins (Arc^20,21^, PEG10^4^) serving as positive controls. A t-SNE visualization confirmed that the 17 selected candidates adequately represent the diversity of the full candidate set (Fig.3f), supporting their use in downstream experimental studies. Notably, one previously established nanocage□forming protein, PNMA2^20,21^, did not pass the dual□model screening strategy, as it was identified only by Model□2. For comparison, PNMA2 was included as a positive control alongside Arc and PEG10.

### Experimental Validation of Candidate Nanocage-forming Proteins

Building on our computational predictions, we experimentally assessed 18 candidate proteins (including PNMA2) for their ability to form mRNA-delivering nanocages. Two previously characterized proteins, Arc and PEG10, both of which contain native RNA-binding domains, served as positive controls for RNA encapsulation efficiency^4,20,21^. In contrast, none of the remaining 16 candidates exhibited recognizable RNA-binding domains when aligned against a curated nucleocapsid protein database using Diamond BLAST (Supplementary Fig.11). To standardize mRNA packaging capability across all candidates, we adopted a modular engineering approach by fusing a C-terminal MS2 coat protein domain, an N-terminal PLCD domain, and an HIV-1 P6 secretion signal peptide (Fig.4a and Supplementary Fig.12)^17^. This design enabled consistent assembly and functional evaluation.

To assess nanocage formation, HEK293T cells were co-transfected with three constructs: (i) the engineered packaging plasmid, (ii) a VSV-G envelope plasmid for pseudotyping to enhance cellular uptake and (iii) a reporter plasmid encoding Cre-NLS mRNA flanked by MS2 stem loops for specific binding^17^ (Fig.4a). After 72 hours, secreted particles were purified from the supernatant and analyzed by transmission electron microscopy (TEM) to visualize capsid-like structures, and by a Cre-loxP reporter assay^4^ to assess functional mRNA delivery (Fig.4b). In the reporter assay, successful delivery of Cre-NLS mRNA induces site-specific recombination and activates EGFP expression in HEK293T cells (Fig. 4c and Supplementary Fig.13).

**Fig. 4.**
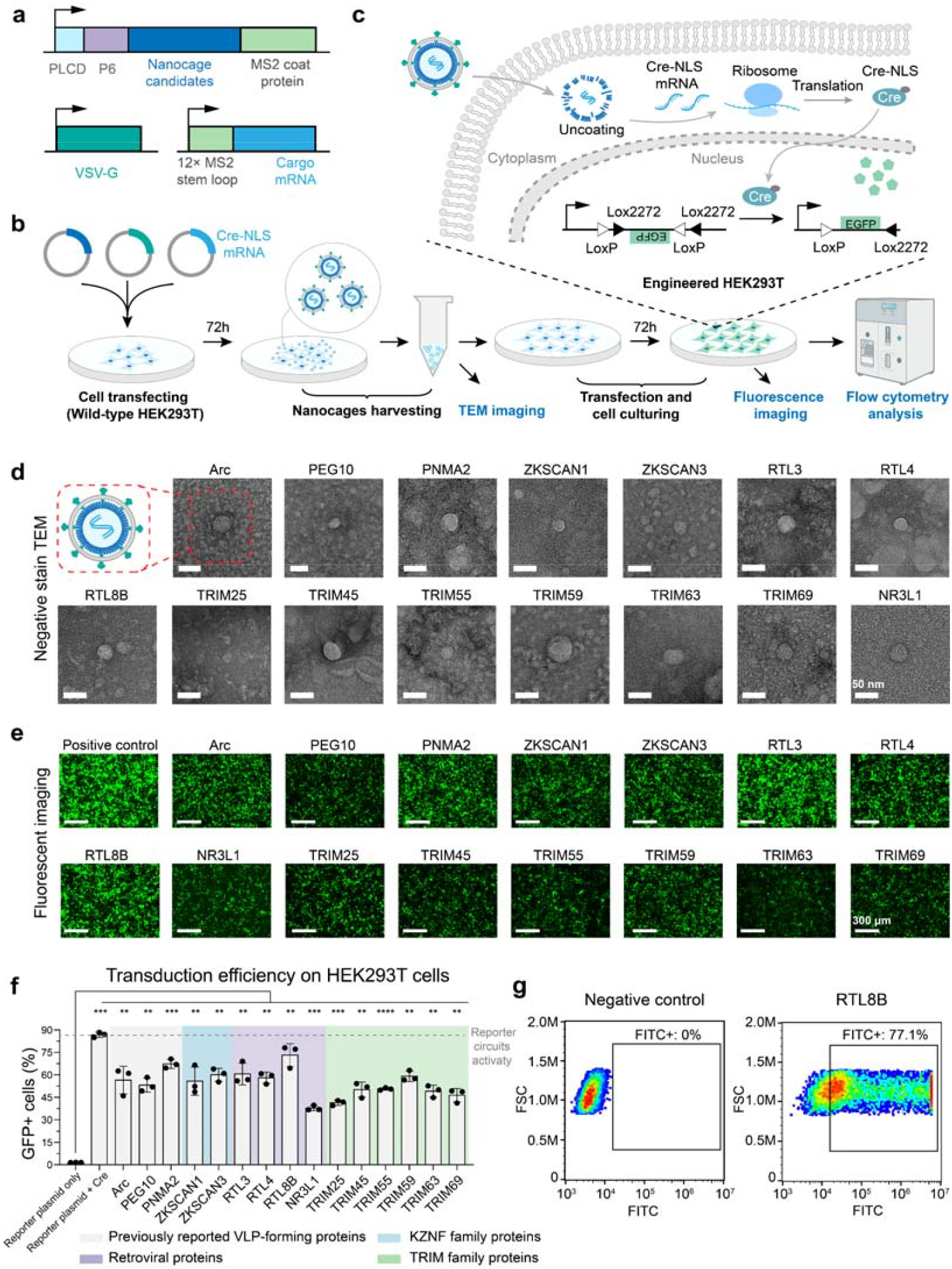
Experimental validation of candidate nanocage-forming proteins for mRNA delivery. **a**, Plasmid design for nanocage production. The packaging plasmid encodes candidate proteins fused to an N-terminal PLCD membrane-binding domain, a C-terminal P6 secretion signal, and an MS2 coat protein RNA-binding module. The fusogen plasmid expresses VSV-G, and the cargo plasmid contains mRNA with 12× MS2 stem loops. **b**, Production and functional testing workflow. HEK293T cells were co-transfected with the three plasmids. Culture supernatants were collected after 72 hours, and nanocages were purified via centrifugation for TEM imaging. Functional validation was performed by transducing Cre-responsive reporter cells and assessing EGFP expression. **c**, Cre-recombinase reporter circuit. Delivery of Cre-NLS mRNA triggers recombination and EGFP expression. **d**, Negative-stain TEM images of purified nanocages. Scale bar, 50 nm. **e**, Fluorescence microscopy of transduced reporter cells. Groups: positive control (Cre plasmid transfection), and nanocages from candidate proteins. Scale bar, 300 μm. **f**, Transduction efficiency quantified by flow cytometry. Data represent mean percentage of GFP-positive cells (*n* =□3 independent experiments, mean ± SD). Statistics by unpaired two-tailed Student’s t-test; P≥0.05, not significant, *P□<□0.05, **P□<□0.01, ***P□<□0.001, ****P<0.0001. From left: reference controls, previously reported VLPs (Arc, PEG10, PNMA2), and nanocages from KZNF, retroviral, and TRIM family candidates. **g**, Representative flow cytometry plots for PBS negative control and RTL8B nanocage-treated samples.

Of the 18 candidates, 15 formed regular, capsid-like structures (Fig.4d). While RTL8C, PGBD1, and RN186 failed to assemble, suggesting limitations in the current predictive model for certain self-assembly determinants. All 15 structurally confirmed candidates mediated Cre-dependent EGFP activation, confirming functional mRNA delivery (Fig.4e and Supplementary Fig.14). Flow cytometry revealed that RTL8B achieved the highest transduction efficiency (73.2%), outperforming established reference proteins Arc (56.5%), PEG10 (53.1%), and PNMA2 (67.4%). Candidates such as ZKSCAN1, ZKSCAN3, RTL3, RTL4, and TRIM59 performed comparably to Arc and PEG10, while other TRIM family members and NR3L1 showed moderate efficiency (Fig.4f). All values were normalized against a PBS-treated negative control (Fig.4g and Supplementary Fig.15).

To assess the safety of the obtained human endogenous nanocages for mRNA delivery, we compared their innate immunogenicity with conventional transfection methods. When applied to phorbol 12-myristate 13-acetate (PMA)-differentiated THP-1 macrophage-like cells, nanocage formulations triggered minimal inflammatory responses, with TNF-α and IL-1β secretion markedly lower than that induced by Lipofectamine 3000–mediated plasmid transfection, reduced to approximately 1/122 and 1/44, respectively. These responses were also comparable to those of untreated controls, indicating that nanocage exposure alone did not provoke detectable inflammation (Supplementary Fig. 16a-c).

These results demonstrate the efficacy of integrating deep learning-based screening with experimental validation for discovering novel and biocompatible nanocage-forming proteins. Among the 15 newly identified candidates (excluding the known Arc, PEG10, and PNMA2), 12 of them (80.0%) successfully assembled into well-defined particles, and all mediated functional mRNA delivery (Supplementary Table 2). Sequence similarity analysis using Diamond BLASTP revealed low homology to previously reported nanocage-forming proteins. Only TRIM55 and TRIM63 showed 40-50% similarity, while TRIM25, TRIM45, and TRIM59 had 0% similarity, and the remainder fell below 40% (Supplementary Fig.17). This indicates that conventional homology-based methods would have missed most candidates, underscoring the advantage of an AI-driven discovery pipeline for identifying structurally capable yet sequence-divergent human proteins suitable for nanocage engineering.

### In-depth Analysis of TRIM Proteins as Nanocage-Forming Candidates

A notable outcome of our experimental validation was the consistent ability of all tested TRIM family proteins to form nanocages and mediate mRNA delivery *in vitro*. This finding is intriguing given the established role of TRIM proteins in antiviral defense^44,45^ and exogenous RNA recognition^46^, prompting inquiry into the compatibility between their innate immune functions and engineered nanocage assembly. For instance, although TRIM25 received a high prediction score from our 1D-CNN model, it exhibited relatively low infectivity (41.3%), indicating a gap between in-silico prediction and delivery efficiency in reality.

To elucidate the mechanistic basis of this discrepancy, we performed a multi-faceted analysis integrating phylogenetics, protein representation learning, and interpretable AI. Protein embeddings generated by the 1D-CNN model revealed distinct clustering of TRIM sequences (Supplementary Fig.18), indicating that the model captures structural and functional heterogeneity within this family. Phylogenetic mapping of Swiss-Prot-annotated TRIM proteins, overlaid with model predictions, further refined this landscape (Fig.5a-b, and Supplementary Fig.19): nanocage-positive predictions clustered within subfamilies C-I, II, IV, V, VI, IX, X, and XI, whereas negative predictions were enriched in C-III, VIII, and Unclassified (UC) group. Subfamily C-VII exhibited internal bifurcation, suggesting divergence in domain architecture and assembly potential.

**Fig. 5.**
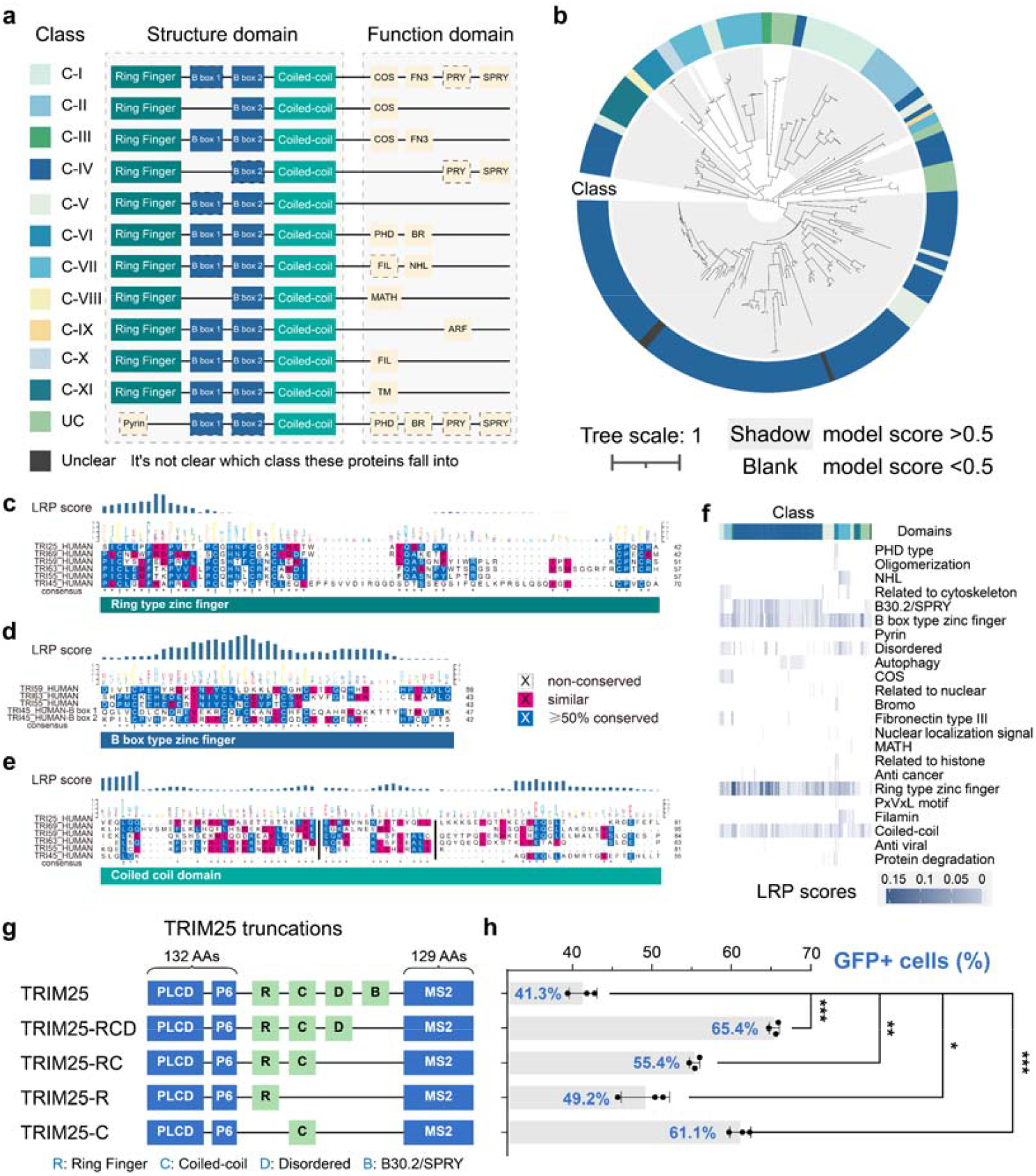
Modular engineering and functional analysis of TRIM25-based nanocages. **a**, Domain architecture of TRIM family proteins. Conserved domains define subfamily. **b**, Phylogenetic tree of TRIM proteins across species, colored by subfamily classification. Branches with model-predicted nanocage propensity scores >0.5 are shaded. **c-e**, LRP score distributions for three common domains. Per-position LRP scores were obtained by first applying min–max normalization to the residue-wise LRP values of each sequence (yielding scores in the range 0–1), and then summing the normalized scores across all sequences at each column of the multiple sequence alignment. **c**, RING-type zinc finger; **d**, B-box zinc finger; and **e**, coiled-coil domain. Higher LRP values indicate stronger model-assigned importance in cage formation. **f**, Heatmap of domain-wise LRP scores across TRIM proteins, clustered by subfamily. **g**, Gene structures of TRIM variants. **h**, Transduction efficiency of modularly engineered TRIM25 variants (*n* =□3 independent experiments, mean ± SD). Significance relative to full-length TRIM25 is indicated. *n* =□3 independent experiments, mean ± SD. Statistics by unpaired two-tailed Student’s t-test; P≥0.05, not significant, *P□<□0.05, **P□<□0.01, ***P□<□0.001.

Since TRIM subfamilies can be characterized by conserved domain combinations^47,48^ (Fig.5a), we hypothesized that specific domain modules govern nanocage functionality. To test this, we applied LRP to identify residues critical for model predictions. Multiple sequence alignments of experimentally validated TRIM constructs (TRIM25, 45, 55, 59, 63, and 69) revealed strong overlap between conserved residues and high-LRP regions (Fig.5c-e), indicating that nanocage assembly is driven by domain-level features rather than isolated motifs. Similar domain-specific patterns were observed in Transformer attention maps, though the 1D-CNN LRP exhibited greater residue-level variation (Supplementary Fig.20).

To systematically evaluate domain modularity, we aggregated residue-level LRP scores across all UniProt-annotated domains in Swiss-Prot-curated TRIM proteins (Supplementary Fig.21), and visualized domain-averaged relevance as a heatmap (Fig.5f). A gradient from gray to dark blue indicated increasing relevance to nanocage formation, while white denoted domain absence. This analysis revealed consistent high relevance for the RBCC module — comprising the RING-type zinc finger, B-box, and coiled-coil domains—whereas domains implicated in unrelated processes (e.g., autophagy) showed low relevance. These findings confirm that domain organization, particularly the RBCC module, is a principal determinant of nanocage assembly, corroborating experimental data highlighting the essential role of the N-terminal RBCC domain (Supplementary Fig.22). Collectively, this modularity framework explains functional heterogeneity within the TRIM family and provides a rational basis for engineering protein-derived nanocages with enhanced functionality.

### Modular Engineering of TRIM Family Proteins for Enhanced Nanocage-based mRNA Delivery

Guided by domain-level relevance mapping, we employed a modular engineering approach to optimize TRIM proteins as mRNA-delivering nanocages. Using TRIM25 as a prototype, we designed a series of truncation mutants to dissect the functional contributions of specific domains to nanocage assembly and delivery efficiency. Domains selected for modification included the RING-type zinc finger (R), implicated in ubiquitination^49^, the coiled-coil domain (C), essential for oligomerization^50,51^, intrinsically disordered regions (D), involved in promiscuous biomolecular interactions^52^, and the B30.2/SPRY domain (B), known for immune recognition and antiviral functions^53^ (Fig.5g). Modifications were designed to preserve core multimerization motifs while reducing potential interference with host immune pathways.

All engineered TRIM25 variants retained the ability to form nanocages and mediate mRNA delivery in HEK293T cells (Supplementary Fig. 23 and 24). Each variant also exhibited consistent improvements in mRNA encapsulation and transduction efficiency compared to full-length TRIM25 (Fig.5h and Supplementary Fig.25). The TRIM25-RCD variant (retaining R, C, and D domains) achieved the highest transduction efficiency (65.4%), a 24.1% increase over wild-type TRIM25 (41.3%). Other constructs also showed enhanced performance: TRIM25-C (61.1%), TRIM25-RC (55.4%), and TRIM25-R (49.2%).

Pairwise domain comparisons revealed distinct functional roles. Removal of the B30.2/SPRY domain markedly improved transduction, consistent with its known role in activating immune pathways^53^; its excision likely mitigates unintended host interactions. Furthermore, comparison between TRIM25-RC and TRIM25-C indicated that removal of the RING domain conferred an additional 5.7% gain in infectivity (*\*\*P*<0.01). Although the RING domain received high relevance scores in our AI model and is critical for hexagonal lattice formation and ubiquitination in TRIM proteins^44,46,49^, its removal may suppress ubiquitin-mediated degradation, thereby enhancing nanocage stability and delivery efficiency.

These results support a domain hierarchy model for TRIM-based nanocages, in which the B30.2/SPRY domain acts as a negative regulator of delivery, the RING domain is necessary for assembly but partially detrimental to function, and an optimal configuration that integrates the RING, coiled-coil, and disordered regions with the PLCD-P6-MS2 cassette maximizes delivery performance. Although improvements were incremental, this study establishes that AI-informed domain editing can systematically enhance the functionality of human protein-derived nanocages for therapeutic mRNA delivery.

## Discussion

By integrating large-scale data curation, machine learning modeling, and experimental validation, we have uncovered a repertoire of human proteins capable of forming virus-like nanocages. Unlike conventional RNA carrier systems that rely on exogenous viral capsids, our approach systematically explores the human proteome, revealing protein families and sequence variants not previously associated with nanocage assembly. This expanded search space underscores the latent structural and functional versatility of endogenous human proteins and provides a rich resource for engineering tailored gene delivery systems suited to diverse therapeutic contexts.

A major strength of our workflow lies in its integration of artificial intelligence with both computational and experimental validation, bridging predictive modeling and biological function for the discovery of protein-based delivery tools. Compared to conventional homology-based methods, our AI-driven pipeline captures long-range dependencies and non-linear feature interactions, enabling the identification of functional candidates with low sequence similarity to known capsids. This approach reduces false positives and reveals proteins overlooked by alignment-based strategies, offering transformative potential for accelerated design and engineering of novel delivery systems.

Furthermore, while deep learning models are often considered “black box” systems^54^, our use of interpretable AI, such as layer-wise relevance propagation, illuminates critical domains and motifs, including various zinc-finger regions (e.g., RING-type, C2H2, SCAN domains), that contribute to self-assembly and nanocage formation. These insights may inform future efforts in rationally engineering capsid proteins to improve cargo loading, enhance delivery efficiency, and reduce immunogenicity^19^.

Several promising directions emerge for future work. First, *in vivo* studies will be essential to evaluate the tissue specificity, pharmacokinetics, and immunogenicity of these novel nanocages. Second, overcoming biological barriers, such as the blood-brain barrier or the tumor microenvironment, could enable targeted applications in gene editing and oncology. Third, domain-level modular engineering, exemplified by our TRIM variants, can be extended to optimize the trade-off between capsid stability and evasion of host immune responses. Fourth, beyond predictive screening, de novo and rational design of nanocage proteins using protein language models holds considerable promise, including applications in structure prediction^55-57^, inverse folding^58^, and conditional sequence generation^35,59,60^. Finally, the generalizable framework presented here, combining systematic data mining with interpretable AI, empirical validation, and AI-informed rational design, can be adapted to other protein engineering challenges, such as discovering membrane fusion proteins or immunomodulatory factors.

In summary, the discovery of numerous human proteins with nanocage-forming potential substantially expands the toolbox for mRNA packaging and delivery. By building a library of human-derived capsid scaffolds, we pave the way for safer, more precise, and personalized delivery systems that minimize off-target effects and immune recognition. The synergy of computational prediction, high-throughput screening, and mechanistic dissection, established in this work, defines a new paradigm for precision nanocage engineering, with broad implications for gene therapy, vaccine development, and regenerative medicine.

## Acknowledgements

We thank George M. Church for advice in this research, Yuting Chen for help in nanocage production and purification, Guanyang Su for help in TEM imaging, and Peilin Zhao for discussion in the early stage of this project. This work was partially sponsored by the National Key R&D Program of China (2025YFA0923500 to C.Z.), the Open Funding Project of the State Key Laboratory of Biopharmaceutical Preparation and Delivery (No. 2024KF-05 to B.A.), the National Science Fund for Distinguished Young Scholars (32125023 to C.Z.), the National Natural Science Foundation of China (No. 62406295 to L.Q.L, 32201105 to B.A.), the Research Grants Council of the Hong Kong Special Administrative Region, China, (Project T45-401/22-N. to P.A.H), the Project of Digital Cell and Drug Discovery Intelligent Platform at Zhejiang Lab (2022PE0AC04 to P.A.H) and the Shenzhen Science and Technology Program (ZDSYS20220606100606013 to C.Z.).

## Contributions

D.X., L.Q.L., B.A., and C.Z. conceived the concept and directed the research. D.X. designed, synthesized, tested, and characterized all the nanocages. D.X. and Y.L. collected the data for deep learning training in this study, which were further analyzed and processed by L.Q.L. and L.J.L. L.Q.L. and D.X. developed and tested the deep learning models and analyzed the interpretability of the AI models. Q.L., J.Z., K.L., Z.Z., J.Z., and K.L. assisted in cell culture and nanocage production. Z.Y., D.X., and L.Q.L. analyzed both computational and experimental data and prepared the figures. X.W., M.T., and G.C. assisted in manuscript revision. C.Z., P.A.H., L.Q.L., and B.A. supervised the whole project. The manuscript was written by D.X., L.Q.L., B.A., and C.Z., with contributions from all authors.

## Data availability

The protein sequences used in this study are listed in the supplementary materials **Table S1**. The source code is available at https://github.com/LanqingLi1993/DeepDelivery for inference of trained AI models and reproduction of this work.

