## Supplementary Information for "AI-driven discovery and engineering of human endogenous nanocage proteins for mRNA delivery"

**Method details**

- Dataset Curation:
  - V0.1, V0.2 dataset construction
  - V1, V2 dataset construction
  - Training dataset preparation
- Identification of Nanocage-Forming Proteins Based on Deep Learning
  - Feature extraction
  - Model backbone
  - Output head
- Benchmarking AI models and comparisons to Diamond and HMMscan
- Screening dataset construction
- Screening potential capsid from human proteins
- Prediction of RNA-binding domains in candidate proteins
- Plasmids
- Cell culture
- Nanocages production and purification
- Negative stain TEM
- Nanocages transduction
- Fluorescent microscopy imaging of nanocage transduced cells
- Flow cytometry analysis
- THP-1 stimulation and cytokine quantification by ELISA
- Phylogenetic analysis of TRIM family proteins
- Interpretable AI Analysis for Deep Learning Models

**Supplemental figures**

**Supplemental tables**

**Method**

**Dataset Curation**

To enable the identification of human nanocage-forming proteins, at the early stage of the project, we experimented with various strategies for data collection. This results in 3 datasets, which we name V0.1, V0.2, and V1 in chronological order, where the final V1 version was used for all reported results in the main text.

1) **V0.1 dataset construction (Supplementary Fig.1b):** A total of 149,167 transposon-derived protein sequences, which were retrieved from the Dfam database and annotated via Diamond BLASTX sequence alignment against Swiss-Prot. Swiss-Prot entries lacking nanocage annotations were appended as negative examples. However, such annotation procedure was later found to be severely biased.
2) **V0.2 dataset construction (Supplementary Fig.1c): N**atural protein sequences annotated with “capsid” keywords in UniProt were collected as positive examples, with all remaining Swiss-Prot entries used as negative examples. Although V2 partially mitigated annotation biases present in V1, models trained on both datasets exhibited limited generalization performance on unseen sequences, underscoring the need for a more biologically diverse and less noisy dataset.
3) **V1 and V2 dataset construction (Supplementary Fig.1a):** A more refined curation strategy was implemented using the InterPro database, which integrates functional annotations from Pfam, TIGRFAMs, and CATH-Gene3D. Keyword-based queries (“capsid”, “viral coat”, “viral capsid”) were applied to identify virus-like particle (VLP)-associated protein families. This search yielded 472 candidate InterPro entries. Each entry was manually reviewed, and families explicitly annotated as capsid proteins or other self-assembling proteins (e.g., IPR000574: Tymovirus coat protein; IPR003309: SCAN domain, similar to HIV capsid domain) were retained. Entries unrelated to VLP assembly (e.g., PF11602: Terminase large subunit) were excluded. The final curated set consisted of 456 high-confidence protein families across 68 viral families, as defined by the ICTV taxonomy, spanning lineages from *Asfarviridae* to *Phenuiviridae*. V2 dataset was a subset of V1 with only retrovirus-derived sequences as positives, with a balanced set of negatives (~1:1 ratio).

**Training dataset preparation:** Two training datasets were prepared for capsid-screening model development.

1）The **complete (V1) dataset** contained 544,311 protein sequences with nanocage-forming potential from the 456 protein families (68 viral families) as positives and 635,089 negative sequences. This dataset was used to train our Model 1 for initial screening **(**Fig.1a**).**

2）A more focused **retrovirus (V2) dataset** contained 251,028 capsid sequences from 22 protein families within *Retroviridae* and 262,322 non-capsid sequences as negatives. As a subset of the complete dataset, it was used to train our Model 2 for further screening (Fig.1a).

Negative sequences were proportionally sampled from all InterPro protein families excluding capsid-associated entries. All sequences were retrieved from InterPro (Supplementary Fig.1a).

**Identification of Nanocage-Forming Proteins Based on Deep Learning**

The training and evaluation of deep learning models were performed based on ImDrug ^1^., a machine learning framework for benchmarking a broad spectrum of deep learning models and algorithms on imbalanced drug discovery datasets. Since the original ImDrug focused on processing molecular data involving chemical compounds and reactions, in this study, we extended its functionalities to protein sequences. As shown in Fig.1a, the whole pipeline effectively breaks down to three modules:

**Feature extraction:**

This module converts raw amino acid sequences into numerical representations as input of the downstream neural network backbones. For light-weight model backbones (top branch of Fig.2a), we employed one-hot encoding for CNN and CNN_RNN, Byte-Pair Encoding (BPE) algorithm ^2^ with a vocabulary of about 4000 frequent subwords from DeepPurpose ^3^ (<https://github.com/kexinhuang12345/DeepPurpose/blob/master/DeepPurpose/ESPF/subword_units_map_uniprot_2000.csv>) for Transformer models, respectively. To support pretrained protein language models like ESM, we directly applied their specialized tokenizer from Hugginface transformers (v4.25.1) package.

**Model backbone:**

We provide our own implementation of the light-weight neural networks in the released code base (<https://github.com/LanqingLi1993/DeepDelivery>/lib/backbone/sequence.py). Specifically, our 1D-CNN consists of three convolutional layers, with 4, 8, 12 kernels and 32, 64, 96 filters respectively, followed by a ReLU activation, an average pooling layer and a dense layer to generate a 256-dimensional output vector. We used Huggingface transformers (v4.25.1) package for pretrained protein language model backbones like ESM.

**Output head:**

We used a default MLP classification head, with 3 hidden layers of dimension 256, 128 and 64 respectively, each followed by a ReLU activation. The last layer is a dense layer of output dimension 2 for binary classification.

**Benchmarking AI models and comparisons to Diamond and HMMscan**

To perform AI model selection (Fig.2c-h), we created an 8:1:1 training/validation/test split of the complete V3 dataset to train all models for 100 epochs. The best checkpoint of each trial was selected based on the validation performance. As a sanity check, for each distinct AI model, we repeated the same procedure across three random seeds and observed very consistent results. Each trial was run on a single Tesla V100 GPU on the high-performance computing platform at Zhejiang Lab, and also tested extensively on individual workstations with NVIDIA RTX 3090 GPUs, resulting in no statistically significant discrepancies. All deep learning models were trained with the identical binary cross-entropy loss. Since our datasets are roughly balanced, no additional techniques for handling data imbalance from ImDrug were used.

To demonstrate the power of our proposed AI-driven strategy of nanocage protein discovery, we assessed the performance of deep learning models against two representative bioinformatic tools: Diamond v2.1.11 ^4^ (with e-value threshold of 1E-3) and HMMscan v3.2.2 ^5^ (with score ≥ 30 and aligned fraction ≥ 0.7 as referenced from ^6^), in light of previous studies ^7,8^. To ensure an equal footing for comparison, we query sequences from the test set against the positive samples of the training set as database, and each sequence exceeding the pre-specified threshold is regarded as a predicted positive sample (Fig.2c-h). To create the more challenging dataset of low-similarity sequences (Fig.2i**)**, we clustered the entire V3 dataset using Diamond and randomly split the clusters by the same 8:1:1 ratio. Note that this procedure was done separately for the positive and negative samples. Among annotated positive samples, True positive (TP) is defined as the number of samples that are predicted as positive (nanocage-forming), and False negative (FN) accounts for the rest that are predicted as negative. Similarly, among annotated negative samples, True negative (TN) is defined as the number of samples that are predicted as negative, and False positive (FP) accounts for the rest. We employed six evaluation metrics (Fig.2c-h) for comparison of AI models along with Diamond and HMMscan in terms of binary classification performance, all of which are explicitly defined via TP, FN, TN and FP as follow:

$$\text{Precision Rate}= \frac{TP}{TP+FP}$$

$$\text{Recall Rate}= \frac{TP}{TP+FN}$$

$$\text{False Positive Rate}= \frac{FP}{FP+TN}$$

$$\text{Accuracy}= \frac{TP+TN}{TP+FP+ TN+FN}$$

$$\text{F1 Score}= \frac{2TP}{2TP+FP+ FN}$$

$$\text{MCC}= \frac{TP\times TN-FP\times FN}{\sqrt{(TP+FP)(TP+FN)(TN+FP)(TN+FN)}}$$

For computational time estimation, we calculated the time of processing a single sequence averaged over all query samples (positive plus negative). Diamond and HMMscan were executed on a server comprised of 64 cores (vCPU) and 512 GiB memory, and AI models were run on a single Tesla V100 GPU on the same server.

**Screening dataset construction:** For large-scale screening of human proteins, 204,052 protein sequences (Taxonomy ID: 9606) were downloaded from UniProt. Sequences were filtered to retain those within a length range of 100–1,024 amino acids, resulting in 109,375 entries. In addition, tissue- and organ-specific protein sequences were obtained from the Human Protein Atlas for downstream analysis.

**Screening potential capsid from human proteins**

To identify potential endogenous nanocage-forming proteins from the human proteome, we applied the selected 1D-CNN capsid screening model in combination with a retrovirus-specific model, implementing a dual-model prediction strategy to enhance specificity. Human protein sequences (taxonomy ID: 9606) were retrieved from UniProt (release date: October 12, 2023), resulting in 204,052 entries. Sequences were filtered to retain those with lengths between 100 and 1,024 amino acids, yielding 109,375 entries for screening.

**Model 1** was trained on a comprehensive dataset of capsid-forming proteins from 68 viral families (complete dataset). **Model 2** was trained exclusively on retrovirus-associated capsid proteins and an equal number of randomly sampled non-capsid proteins (1:1 ratio) to mitigate class imbalance. For each sequence, both models generated an independent probability score reflecting the predicted likelihood of capsid or nanocage-forming potential. Only sequences classified as positive by both models were designated as candidates, thus reducing false-positive rates.

Following this initial consensus filtering, we applied a probability score threshold of >0.5 for candidate selection. The resulting hits were visualized via t-SNE based on model embeddings, and sequences were further curated to ensure broad coverage across the t-SNE map by selecting points distributed as evenly as possible. Final candidate sequences were synthesized by GENCEFE Biotech (Wuxi, China).

**Prediction of RNA-binding domains in candidate proteins**

To assess whether human candidate nanocage-forming proteins possess putative RNA-binding regions, we performed sequence similarity searches against a curated reference database of viral and cellular nucleocapsid proteins known to exhibit RNA-binding activity. The reference dataset was compiled from UniProtKB entries annotated with the Gene Ontology term “Nucleocapsid” (GO:0003723) and manually filtered to retain proteins harboring experimentally validated nucleocapsid or capsid RNA-binding domains.

Protein sequences of the candidates selected for experimental validation were queried against the reference database using Diamond BLASTP in sensitive mode, with an e-value cutoff of 1×10⁻⁵ and a minimum sequence identity threshold of 30%. Alignments meeting these criteria and encompassing known RNA-binding motifs (e.g., CCHC-type zinc finger, arginine-rich regions) were considered indicative of potential RNA-binding domains.

**Plasmids**

The sequences of all candidate proteins used in this study are listed in Supplementary Table 1. The envelope plasmid was pMD2.G (Addgene plasmid #12259). Cargo plasmids were constructed based on pFH3.3_Cre‑MS2x12 (Addgene plasmid #205559) and subcloned into the pcDNA3.1 myc‑His A vector (GENCEFE Biotech, Wuxi, China) for expression of reporter mRNAs harboring 12× MS2 stem‑loops. Packaging plasmids were generated by cloning the PLCD, P6, and MS2 coat protein sequences from pFH2.105_PB-EPN24-MCP-T2A-eGFP-PGK-Blast (Addgene, #205550) into the pcDNA3.1 myc-His A backbone and further fused with candidate sequences identified during the screening process.

**Cell culture**

HEK293T cells were incubated at 37℃ with 5% CO2 in complete DMEM medium. Complete DMEM medium consists of DMEM medium (Gibco, 11965092), which was supplemented with 10% fetal bovine serum (Gibco, 10091148), 1% penicillin/streptomycin (Gibco, 15140122), and 1% GlutaMAX™ (Thermo Fisher, 35050061).

**Nanocages production and purification**

HEK293T cells were seeded at a density of 1.2 × 10^7^ cells per 100 mm culture dish (Corning, 430167) and cultured in 6 mL complete DMEM medium, incubating at 37°C with 5% CO₂. After 24 hours, or when cells reached approximately 90% confluence, the medium was replaced with 6 mL Opti-MEM (Gibco, 31985070). After 2 hours, a total of 30 µg of plasmids (10 µg each of packing, envelope, and cargo plasmids) were transfected into the cells using the TransIT-X2 Dynamic Delivery System (Mirus, MIR 6000), following the manufacturer’s instructions. The medium was replaced with 20 mL complete DMEM medium after 6 hours. Cells were incubated for 72 hours.

At 72 hours post-plasmid transfection, the cell culture supernatant was collected and clarified by low-speed centrifugation at 1000 × g for 10 minutes to remove cell debris. The resulting supernatant was then filtered through a 0.45 μm sterile Durapore PVDF membrane filter (Millex-HV, SLHV033RB). The filtrate was then mixed with one-third of its volume of Lenti-X Concentrator (TAKARA, 631232) and then incubated at 4°C overnight. After incubation, the solution was centrifuged at 1500 × g, 4°C for 45 minutes. The supernatant was discarded, and the resulting pellet was resuspended in 0.2 mL complete DMEM medium.

**Negative stain TEM**

Negative stain TEM experiment was performed by Songshan Lake Materials Laboratory with JEOL/JEM-F200 transmission electron microscopy. In sample preparation, 10 μL sample was dropped on a copper grid coated with a continuous carbon film, removed excess solution after 60 seconds by touching the grid with filter paper strips, dropped and removed 10 μL ddH_2_O twice for washing, then 10uL of negative staining solution (uranyl acetate, 1% aqueous solution) was dropped on the copper grid and removed after 60 seconds. Finally, dried the copper grid at room temperature.

**Nanocages transduction**

HEK293T cells were seeded at a density of 2 × 10^5^ cells per well in a 24-well cell culture plate (Corning, 3524) and cultured in 0.5 mL of complete DMEM medium at 37°C with 5% CO₂. After 24 hours or when the cells reached approximately 90% confluence, the medium was replaced with 0.5 mL Opti-MEM (Gibco, 31985070). After 2 hours, 0.5 mg reporter plasmids were transfected into the cells using the TransIT-X2 Dynamic Delivery System (Mirus, MIR 6000), according to the manufacturer’s instructions. The medium was replaced with 0.8 mL complete DMEM medium after 6 hours. The harvested nanocages were resuspended in complete DMEM medium. The nanocage solution was then added to the cell culture medium, with 0.8 μL of Polybrene (Glpbio, GC19206) per 1 mL of medium. The cells were incubated for 72 hours before further use.

**Fluorescent microscopy imaging of nanocage transduced cells**

Fluorescent microscopy imaging of nanocage transduced cells was captured in a 24-well cell culture plate (Corning, 3524). Before imaging, the medium was replaced with 0.5 mL Phosphate-buffered saline (PBS, VivaCell biosciences, C3580-0500). All fluorescent images were captured by ThermoFisher AMF5000. Imaging condition in dark room: GFP channel, light 16.21, exposure 36.6, gain 25.2; TRANS channel, light 56.63, exposure 17.4, gain 3.56.

**Flow cytometry analysis**

Nanocage-transduced cells were cultured in 24-well plates and washed with PBS (VivaCell biosciences, C3580-0500) to remove residual medium. The cells were then treated with 0.5 mL Trypsin-EDTA solution (Gibco, 25300054) for 1 minute to detach the cells. Following digestion, 0.5 mL complete DMEM medium was added to neutralize the trypsin. The cell suspension was centrifuged at 1000 rpm for 3 minutes, and the supernatant was carefully discarded. The cell pellet was resuspended in PBS (VivaCell Biosciences, C3580-0500). The resuspended cells were analyzed using a Beckman CytoFLEX S flow cytometer. Prior to analysis, it is essential to ensure that the cells are fully resuspended. Raw flow cytometry data were exported as FCS files for subsequent analysis.

The FCS files were analyzed using FlowJo v10.9.0 software. Single-cell events were gated using side scatter (SSC-H) and forward scatter (FSC-H) signals to isolate viable cells. Fluorescence intensity was assessed through FSC-H and FITC-H channels. The FITC-positive population was defined by drawing a gating box based on the negative control sample, ensuring the FITC+ rate was set to zero.

**THP-1 stimulation and cytokine quantification by ELISA**
THP-1 cells were seeded in 12-well plates at a density of 2×10^6^ cells per well and differentiated with phorbol 12-myristate 13-acetate (PMA, 50 ng/ml) for 24 h at 37 °C in a humidified incubator with 5% CO2. After differentiation, the supernatant was removed, cells were washed twice with PBS, and fresh RPMI 1640 medium (Gibco, 11875093) was added for a 24-h recovery period. Cells were then incubated for 12 h with virus-like particles (VLPs) in a total volume of 0.5 ml per well (VLP input equivalent to the yield from two 10-cm dishes per well). For contrast, cells were also treated with 200ng/ml lipopolysaccharide and Lipofectamine™ 3000 (Invitrogen, L3000015) with 1.5μg cargo plasmids according to the manufacturer’s instructions. Following stimulation, culture supernatants were collected for cytokine measurements.

TNF-α and IL-1β concentrations in culture supernatants were quantified using a human TNF-α ELISA kit (BOSTER; EK0525) and a human IL-1β ELISA kit (BOSTER; EK0392), respectively, according to the manufacturers’ instructions (Supplementary Fig.16 a).

**Phylogenetic analysis of TRIM family proteins**

We generated multiple sequence alignments iteratively with MAFFT v7.520^9^, removing ambiguously aligned regions using trimAl v1.4^10^. The final alignment was visualized with Texshade ^11^. Phylogenetic trees were inferred from the alignment by maximum likelihood with the LG substitution model implemented in PhyML v3.1^12^, and visualized using the Interactive Tree of Life (iTOL) platform^13^. All software used the default parameter in this part.

**Interpretable AI Analysis for Deep Learning Models**

As neural networks are often considered “black boxes”, to understand how model decision is affected by the input, we employed various strategies to attribute the model output score to each individual amino acid in the protein sequence. Such token-wise scores make the model decision process more interpretable and shed light on the sites/domains of a protein sequence relevant to its nanocage-forming potential, from the AI model perspective.

The major technique employed in this work is Layer-wise Relevance Propagation (LRP), which is widely used in computer vision ^14,15^ and natural science ^16,17^ applications involving modern CNNs. Rooted in the principle of conservation of relevance, LRP works by starting from the network’s final prediction (e.g., the probability of nanocage-forming potential) and recursively propagating this "relevance" backward through each layer of the network—from the output layer down to the input layer. During propagation, it uses layer-specific rules to redistribute relevance scores between neurons, ensuring that the sum of relevance scores at any layer equals the sum of scores at the layer above (preserving total relevance). An illustration of the LRP procedure is provided in Supplementary Fig. 26. The basic propagation rule of the relevance for the neurons is LRP-0:

$$LRP-0: R_{j}= \sum_{k} \frac{a_{j}w_{jk}}{\sum_{0,j} a_{j}w_{jk}}R_{k} ,$$

where $j$ and $k$ represent neurons at two consecutive layers, $R$ is the relevance, $a$denotes lower layer activations and$w$ is the model weight. In practice, to improve the output quality, two enhanced rules are often considered:

$$LRP-\epsilon: R_{j}= \sum_{k} \frac{a_{j}w_{jk}}{\epsilon+\sum_{0,j} a_{j}w_{jk}}R_{k} ,$$

$$LRP-\gamma: R_{j}= \sum_{k} \frac{a_{j}w_{jk}^{+}}{\sum_{0,j} a_{j}(w_{jk}+\gamma w_{jk}^{+})}R_{k} ,$$

where $w^{+}$ is a positive weight, $\epsilon$ and $\gamma$ are dimensionless parameters. In this work, we combined both $LRP-\epsilon$ and $LRP-\gamma$rules and applied $\epsilon, \gamma=0.25$ for all analyses of our 1D-CNN model.

For more complex model backbones such as Transformers, LRP implementation can be challenging due to the overly complicated propagation rules. Fortunately, the attention score naturally operates as a token-level pair-wise relevance score:

$$S_{h}=\text{Softmax} ( \frac{Q_{h}K_{h}^{T}}{\sqrt{D_{h}}}),$$

where $S_{h}$ is the attention score of each head, and $D_{h}$ is the head dimension, $Q_{h}$ and $K_{h}$ represent query and key respectively. By averaging across the head dimension, we obtain an aggregated attention matrix $S_{avg}\in\mathbb{R}^{n\times n}$, which reflects how each token attends to every other token, where $n$ denotes the sequence length. To obtain a token-wise relevance score shown in Supplementary Fig.20, we computed the average attention each token receives from all others in the sequence, by averaging along the first dimension of $S_{avg}$.

With the token-level relevance scores of a given protein sequence, high-relevance regions were identified by applying a threshold to the normalized scores (e.g. 0.5/SeqLen for LRP), and contiguous high-relevance segments were extracted. These predicted important regions were then compared with functional domain annotations retrieved from UniProt to evaluate correspondence between predicted model attention and known protein functional sites. UniProt functional domain annotations used in this study included “Coiled-coil”, “Domain [FT]”, “Repeat”, “Region”, “Motif” and “Zinc finger”. For functional domains that appear less frequently have been grouped into one label, see in Supplementary Table 3. Annotations retrieved from UniProt contained information include start position, end position, name and evidence (reference).

To quantitatively assess relevance at the domain level, per-domain LRP scores were computed by averaging the relevance values across all residues belonging to each annotated domain. Scores were further aggregated across proteins to obtain average relevance per domain category. Visualizations were generated in the form of heatmaps combining LRP-based attribution maps with annotated domain boundaries, enabling the comparison of model-inferred important regions with known structural or functional elements.

**Supplemental figures**

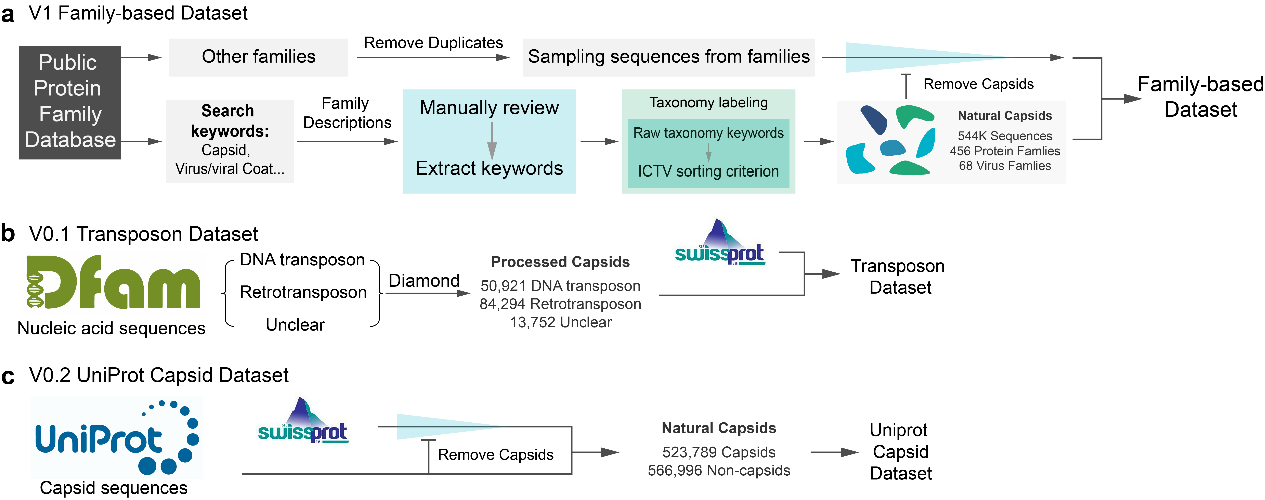

**Supplementary Fig.1.** **Data collection workflows. a**, Schematic of data sourcing and annotation pipeline. Capsid-related entries were retrieved from the InterPro database and manually curated based on family descriptions. Viral taxonomy assignments were standardized using the ICTV classification. The curated capsid sequences were combined with non-capsid entries (excluding duplicates and capsid-related sequences) to generate the final labeled dataset for machine learning. **b-c,** Data collection workflow of the V1 & V2 datasets.

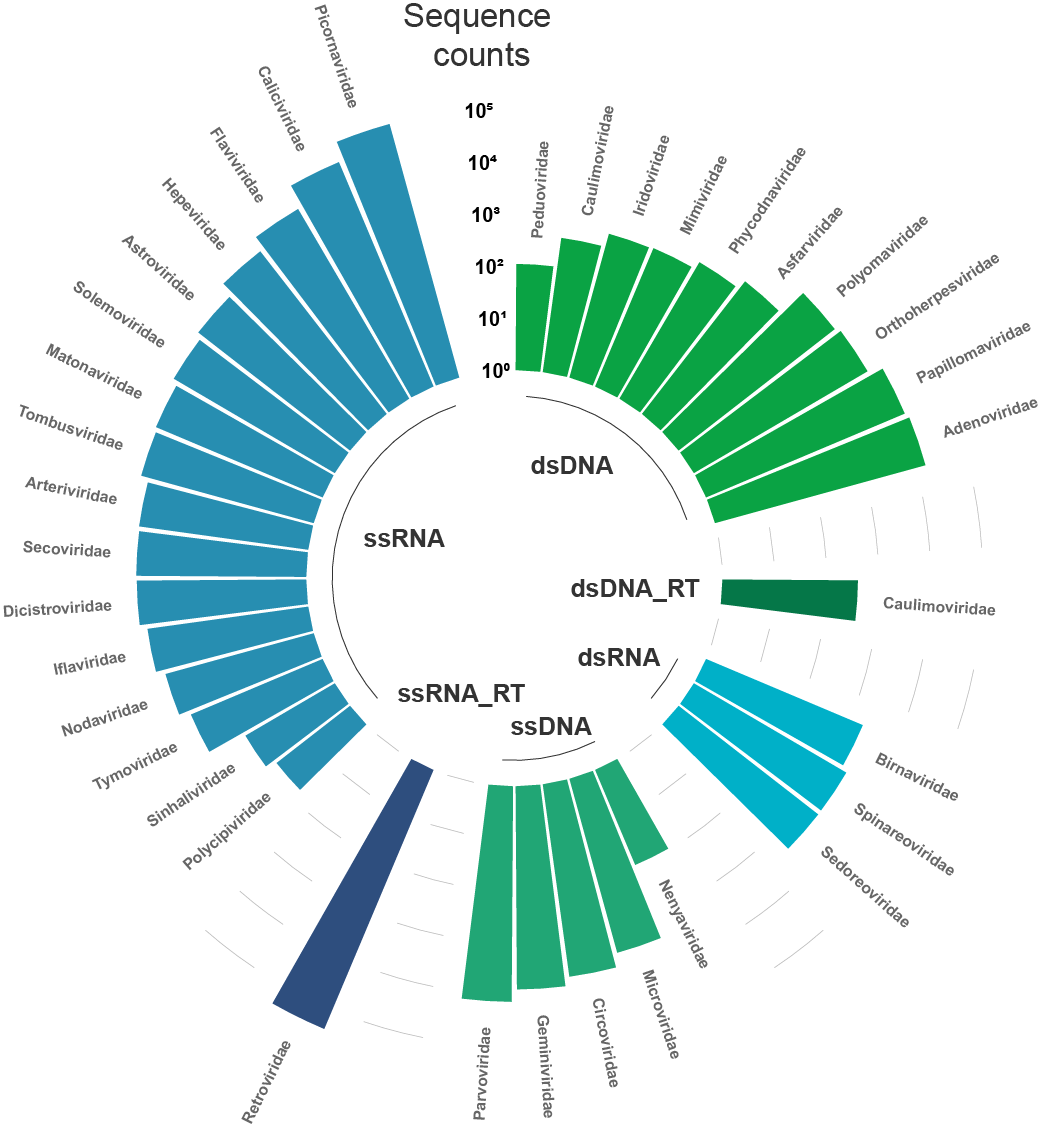

**Supplementary Fig.2.** **Sequence abundance across viral families.** Viral families are grouped by genome type and displayed in a circular layout (Only display families that sequence counts higher than 50). Bar lengths correspond to the number of sequences within each family. This illustrates the high diversity of viral families present in the dataset.

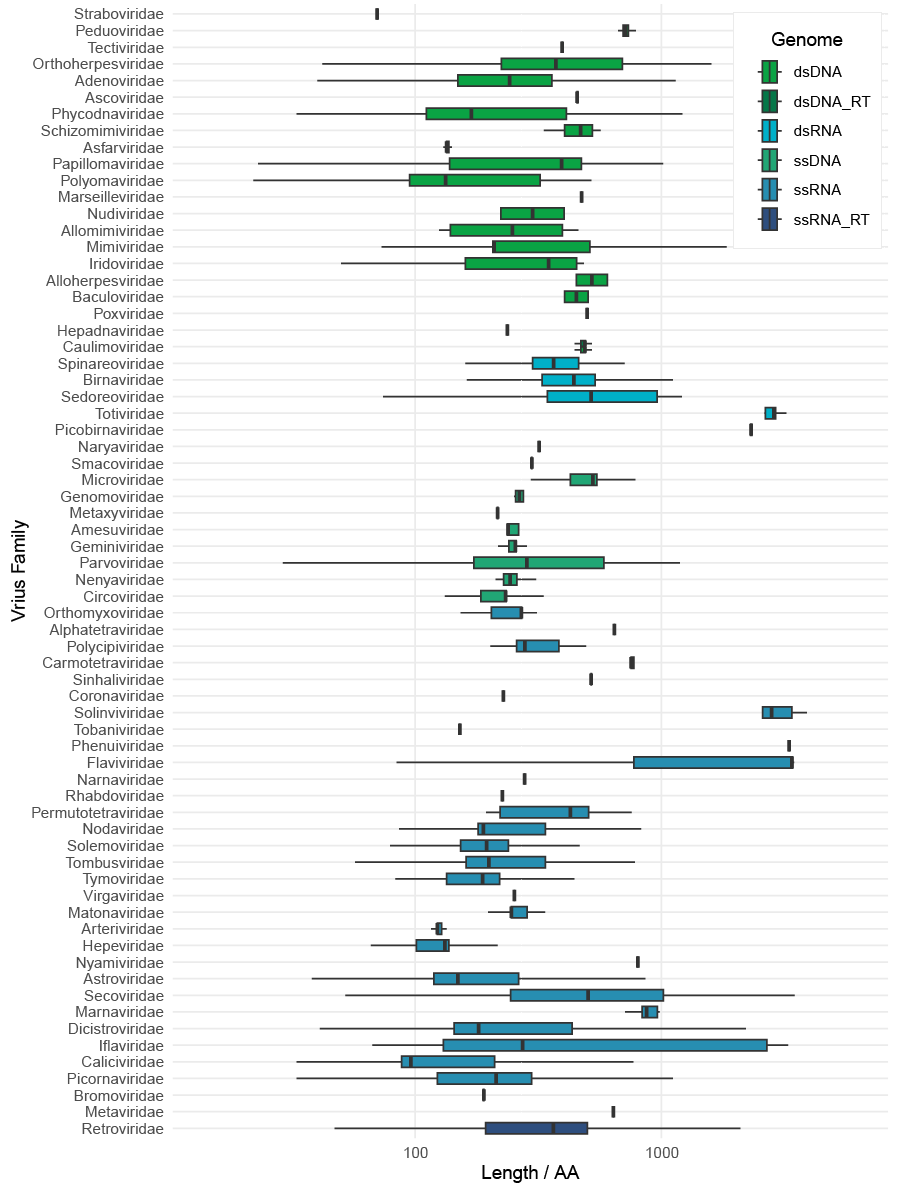

**Supplementary Fig.3. Dataset composition and sequence properties.** Left: Capsid proteins were classified into 68 viral families according to ICTV taxonomy. Right: Sequence length distribution of capsid proteins across viral families. Boxes are color-coded by genome type: light green (dsDNA), green (ssDNA), dark green (dsDNA-RT), light blue (dsRNA), blue (ssRNA), and dark blue (ssRNA-RT). Sequence lengths are plotted on a logarithmic x-axis.

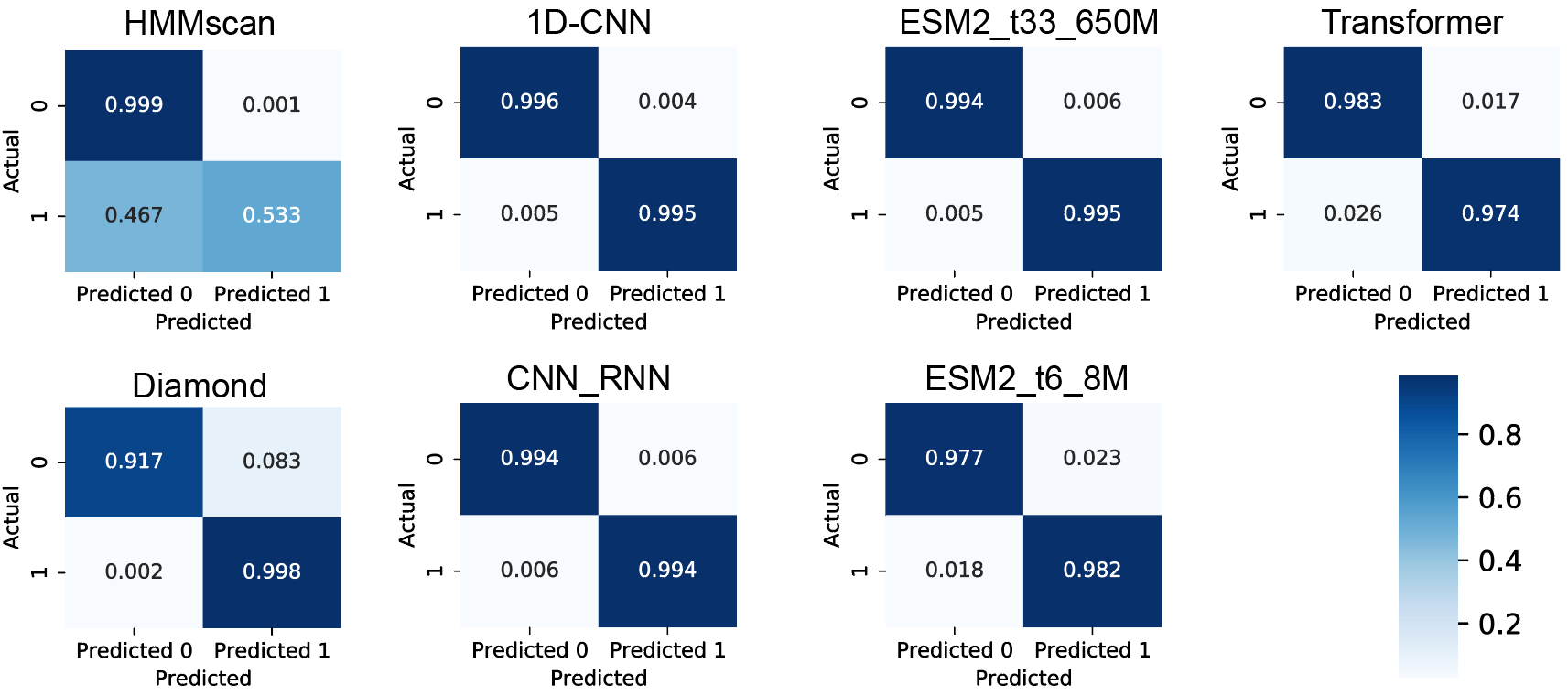

**Supplementary Fig.4. Confusion matrix of bioinformatics methods and deep learning model in the VLP classification task.**

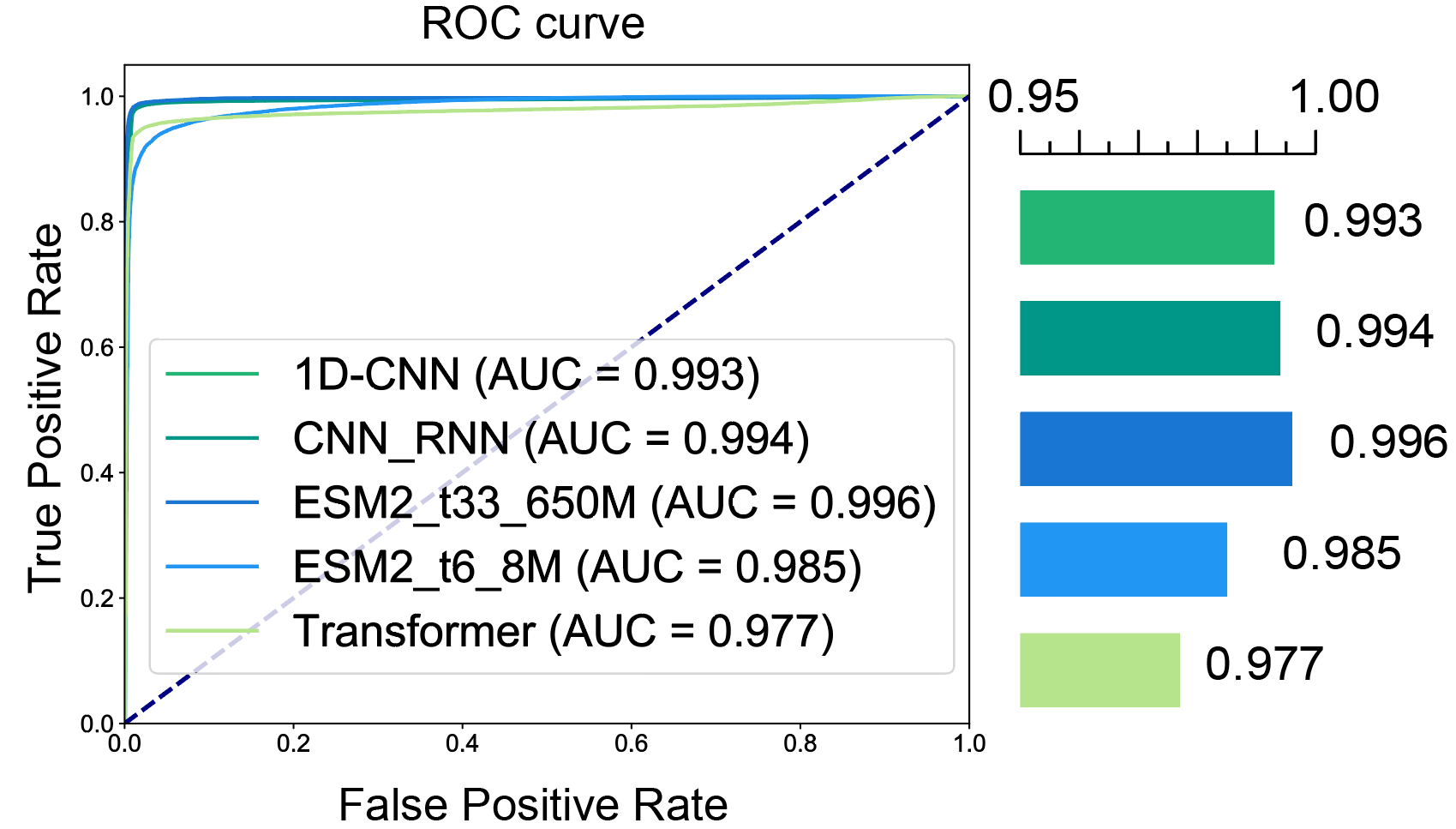

**Supplementary Fig.5. ROC-AUC curve of deep learning models.**

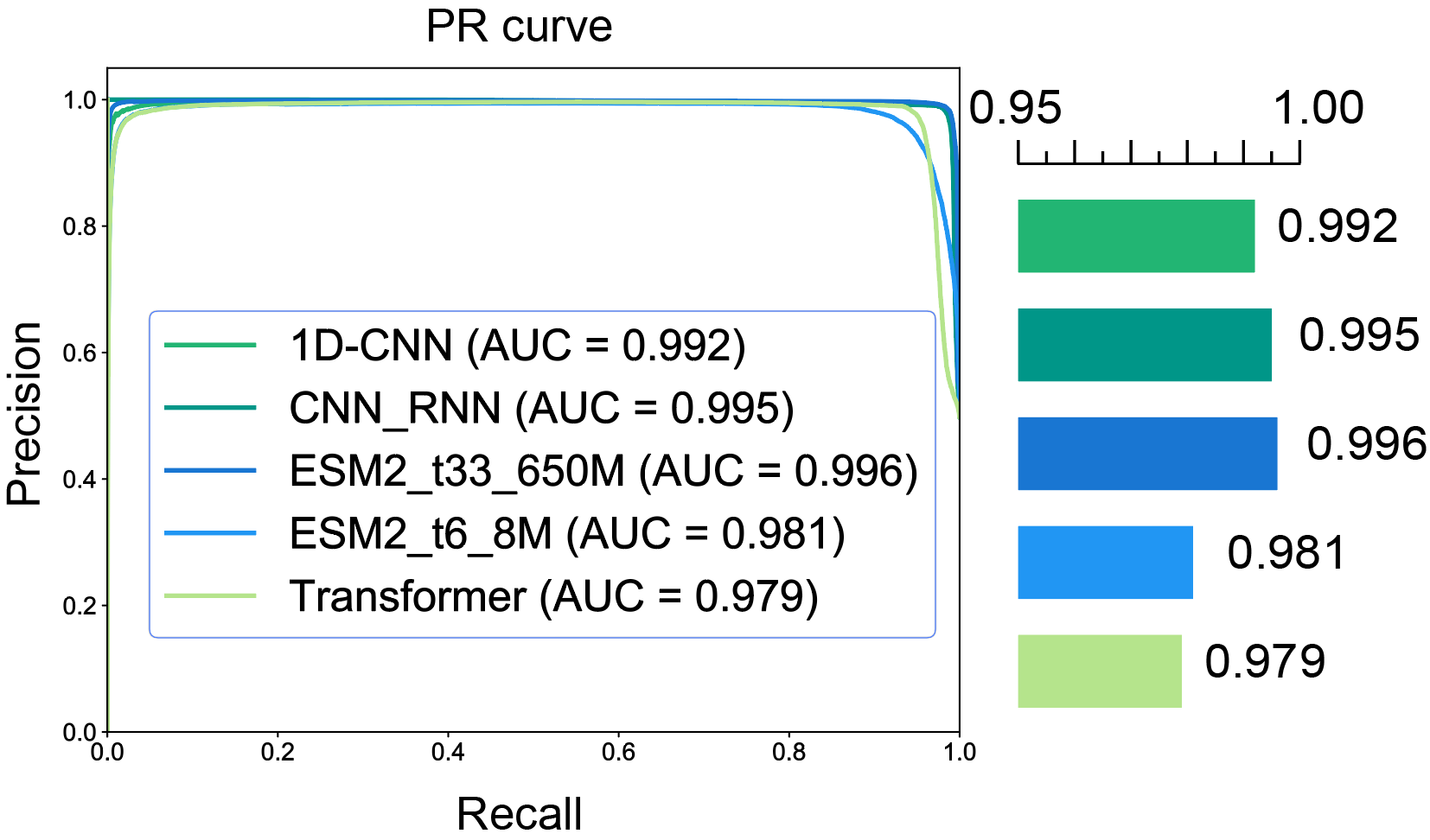

**Supplementary Fig.6. PR-AUC curve of deep learning models.**

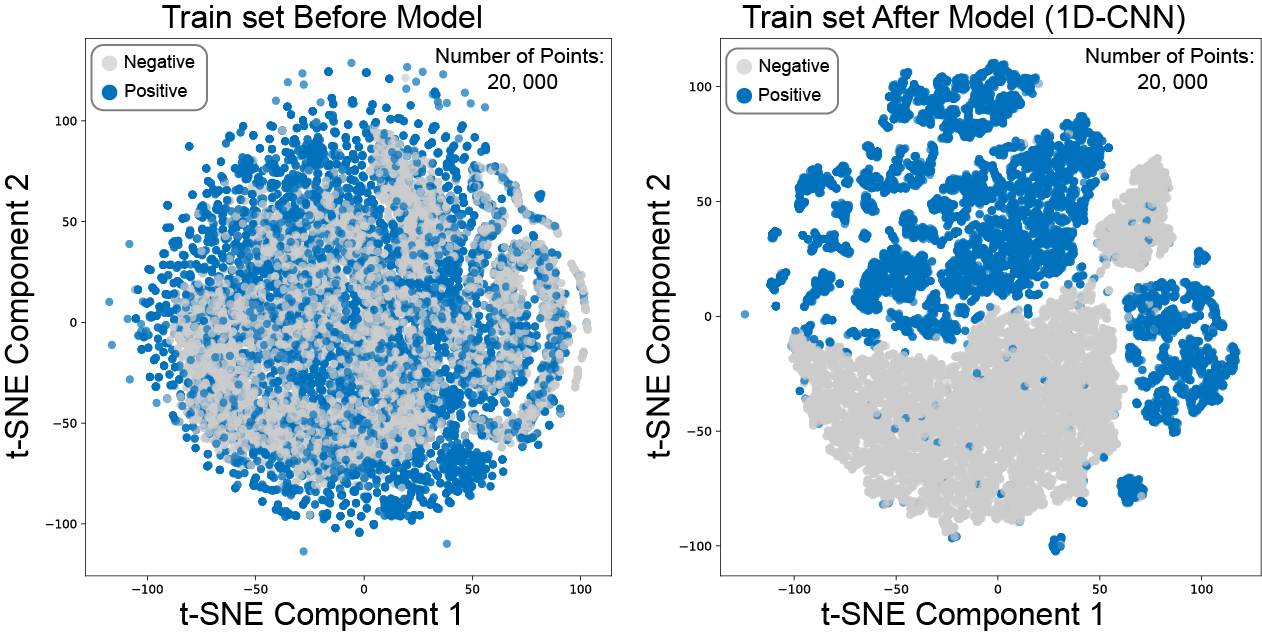

**Supplementary Fig.7.** **T-SNE plot of 20,000 randomly selected sequences in the train set.** Left: Clustered by sequences embedding before the 1D-CNN model. Right: Clustered by sequences embedding after the 1D-CNN model. Gray dots indicated negative samples, while blue dots indicated positive samples.

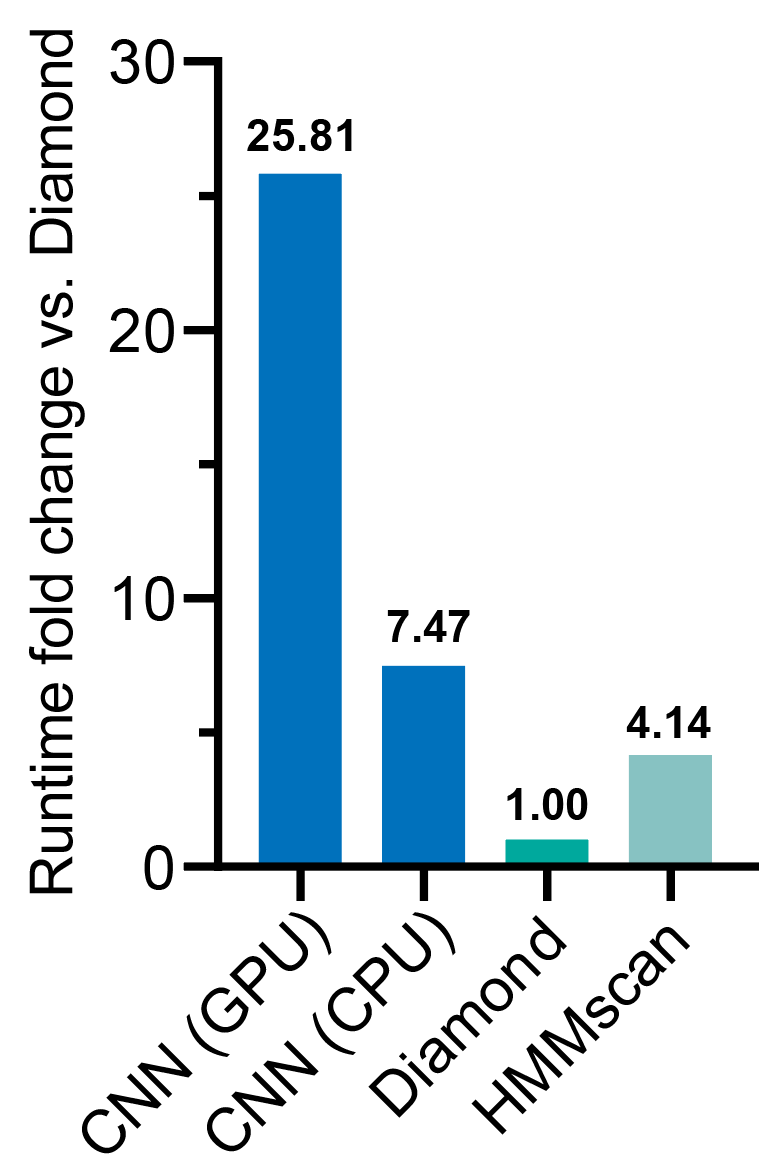

**Supplementary Fig.8.** **Runtime fold change of 1D-CNN and bioinformatics methods on test set.**

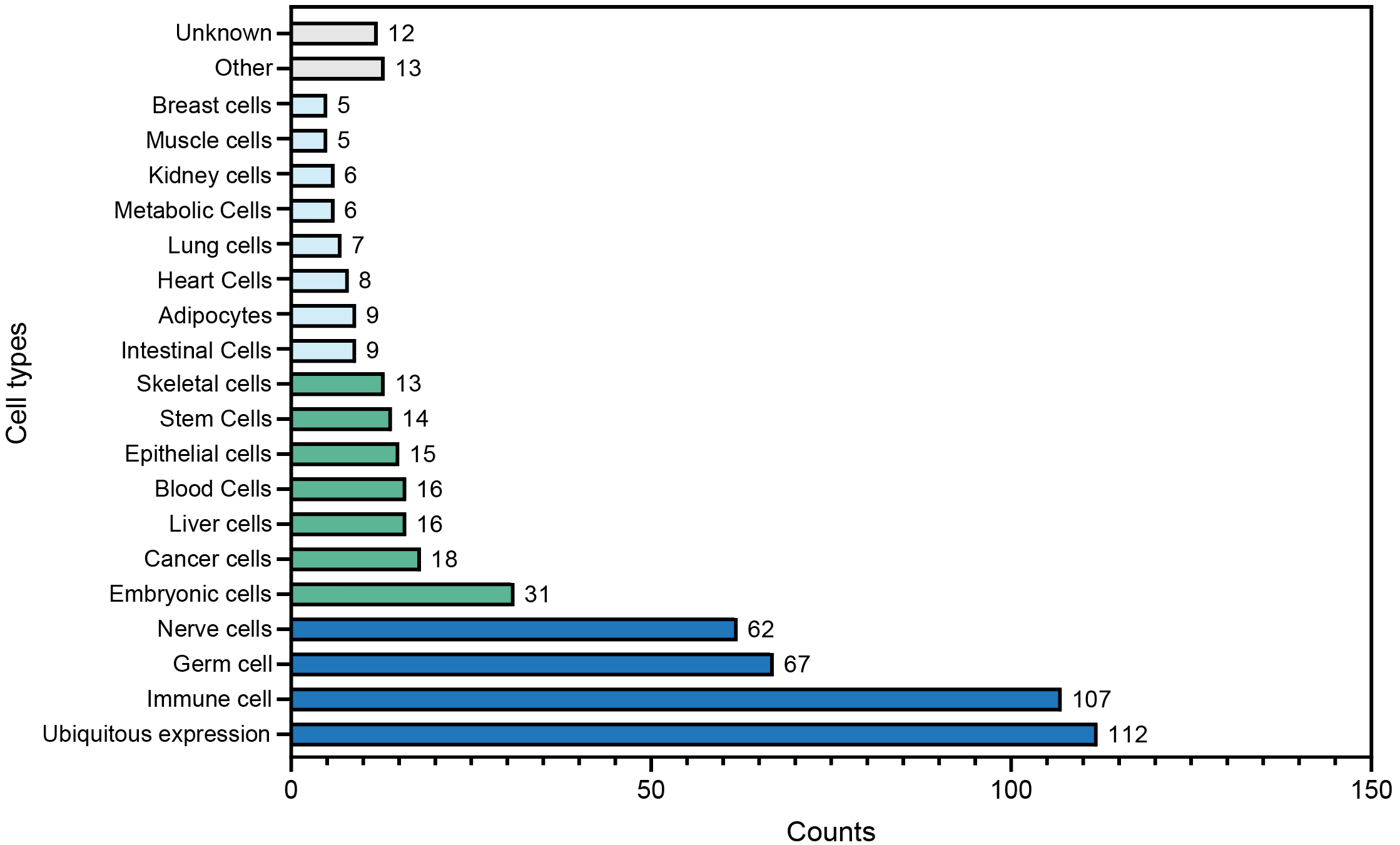

**Supplementary Fig.9. Tissue-specific expression profiles of the 512 candidate proteins.** A single candidate may be expressed in multiple cell types. The number of surfaces in different color ranges falls within different intervals.

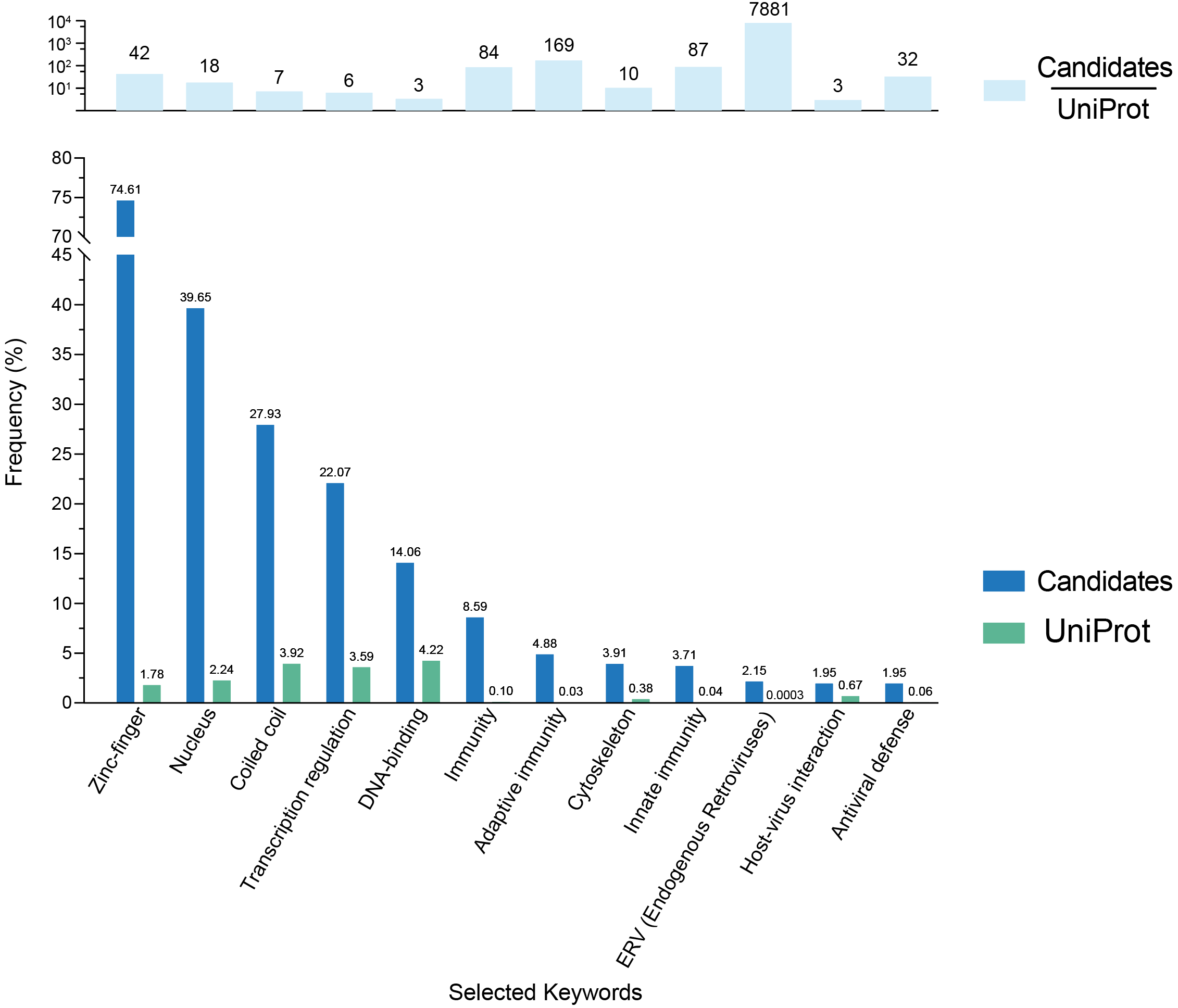

**Supplementary Fig.10.** **Frequency of selected keywords in the 512 candidate proteins and the UniProt database.** Top: Ratio of keyword frequencies between the two datasets. The selected keywords are those related to nucleic acids, immunity, or viruses (including human endogenous viruses). Bottom: Frequency of selected keywords in the 512 candidates and the UniProt database. These results demonstrate that our model enriches keywords potentially associated with viral capsids.

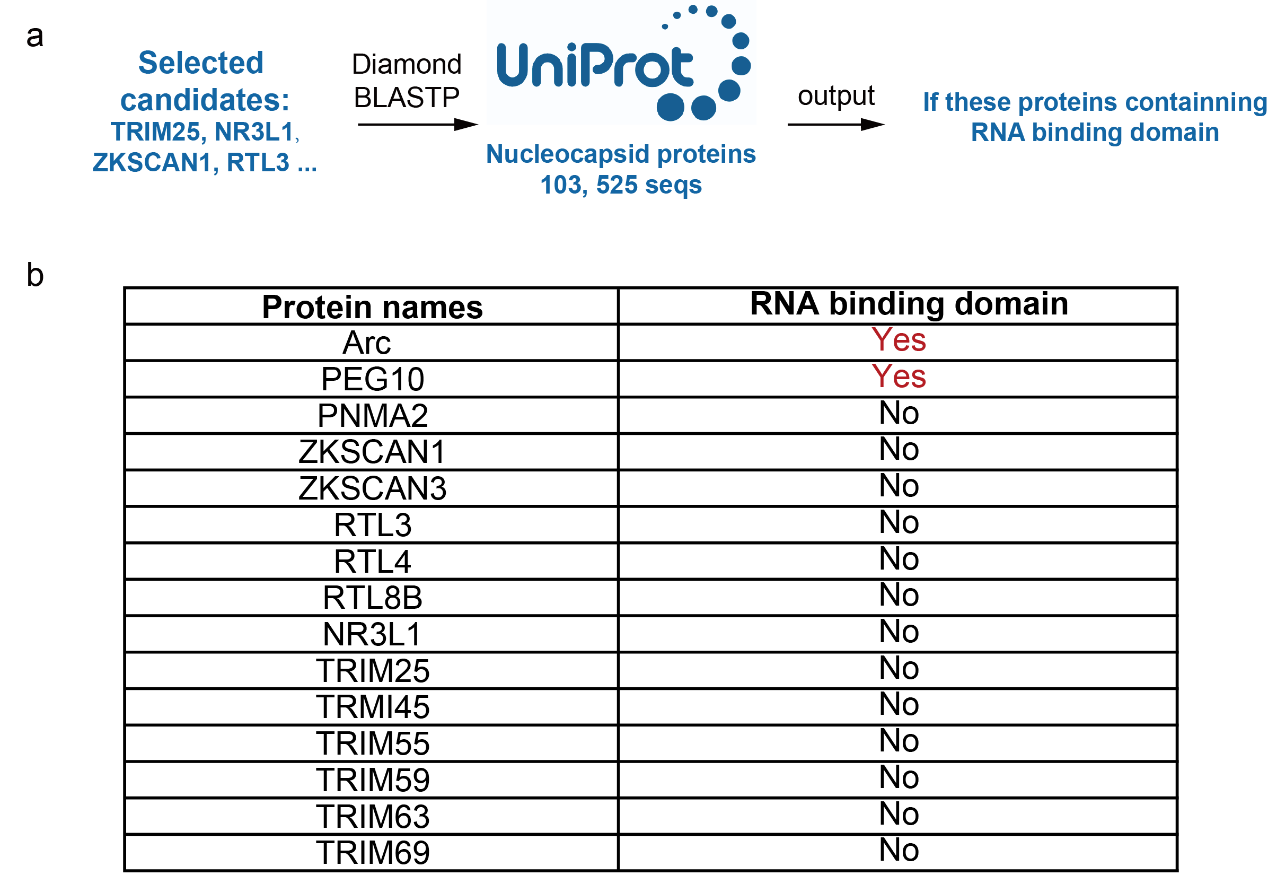

**Supplementary Fig.11. Prediction of RNA-binding domain in selected candidates. a**, RNA-binding domains were identified when comparing the selected candidates against a database of Uniprot nucleocapsid-containing proteins by Diamond BLASTP. **b**, RNA-binding domain analysis results of selected candidates. Only Arc and PEG10 naturally possess RNA-binding domains, as reported in published literature ^18-20^. PEG10 can also be predicted by Diamond BLASTP, while Arc has an atypical nucleocapsid domain, which cannot be predicted by Diamond BLASTP.

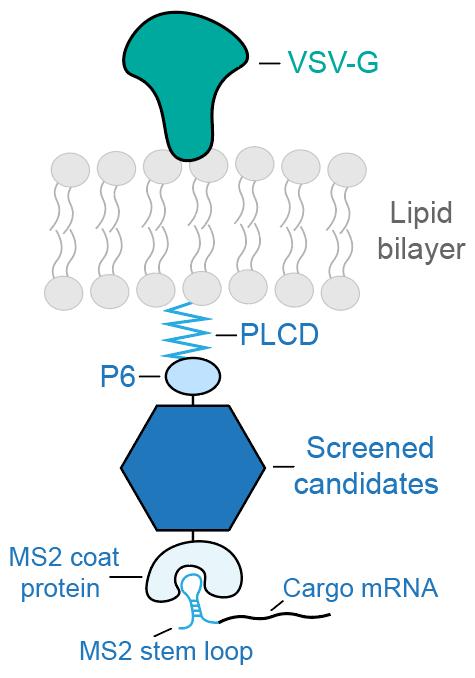

**Supplementary Fig.12. Illustration of a single complex in engineered VLPs.**

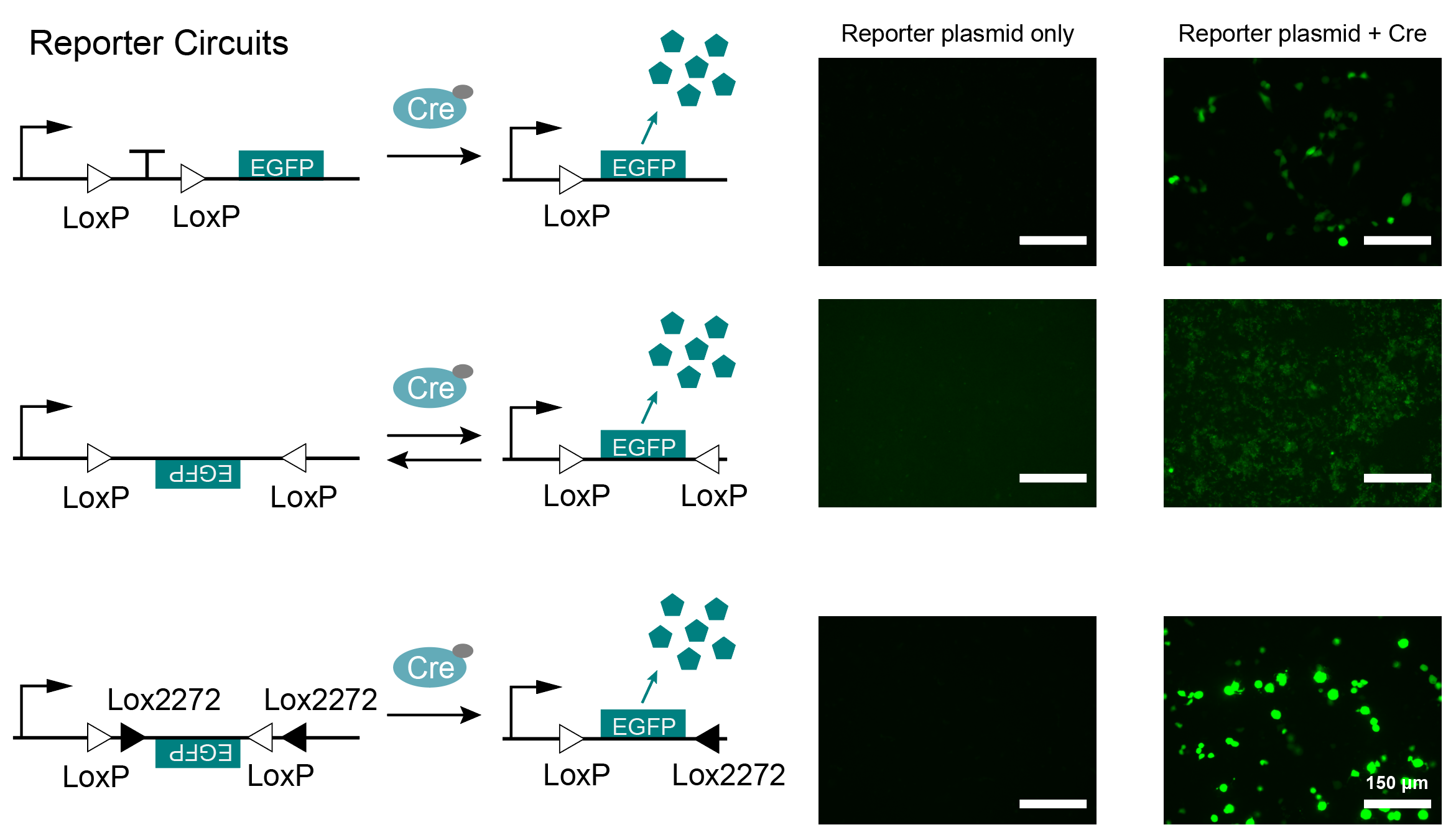

**Supplementary Fig.13. Reporter circuits tested in this study.** Left: illustration of reporter circuits. Right: fluorescent microscope imaging of reporter plasmid only group (negative) and reporter plasmid + Cre group (positive).

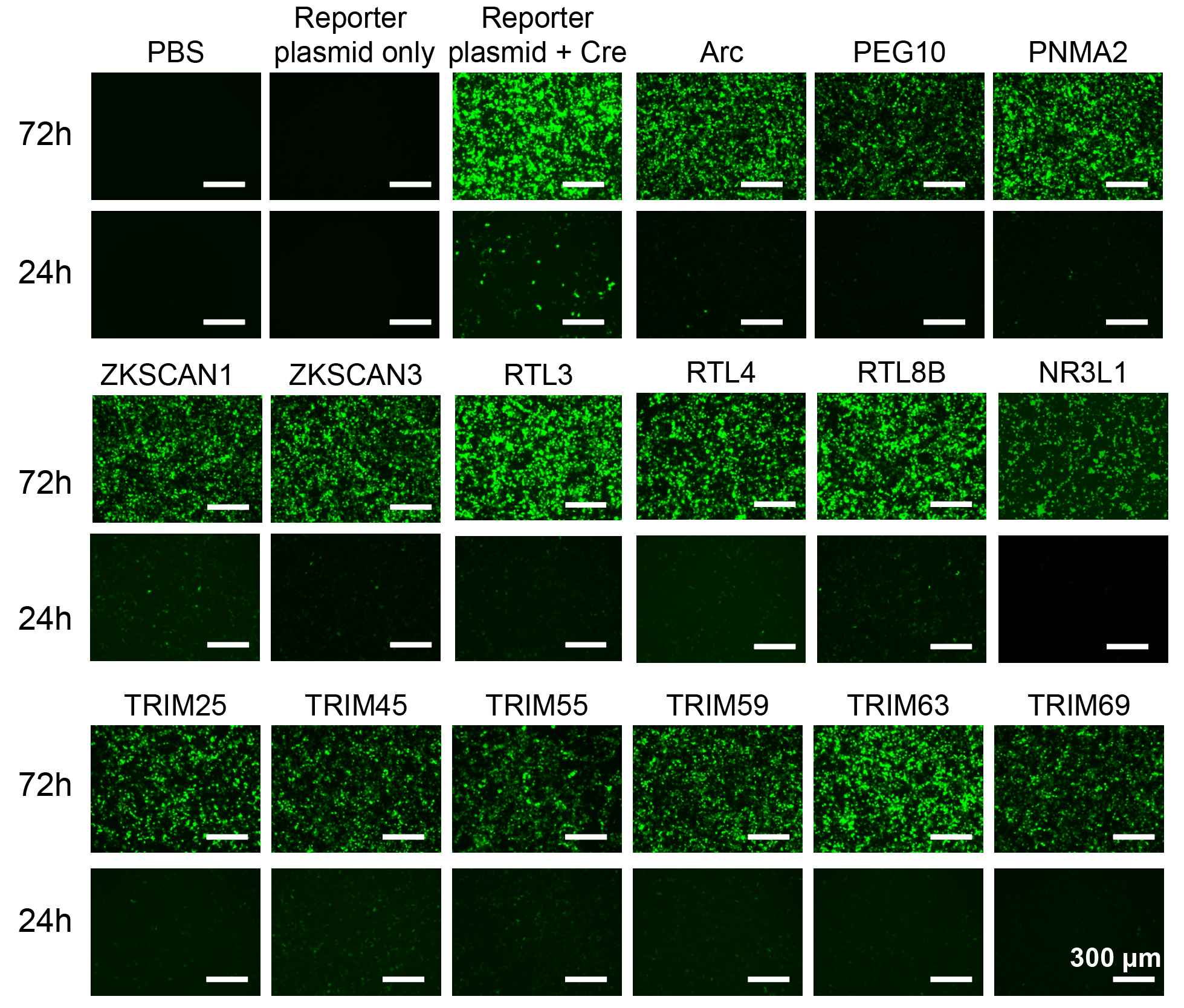

**Supplementary Fig.14.** **24- and 72-hour imaging of transduced HEK293T cells.**

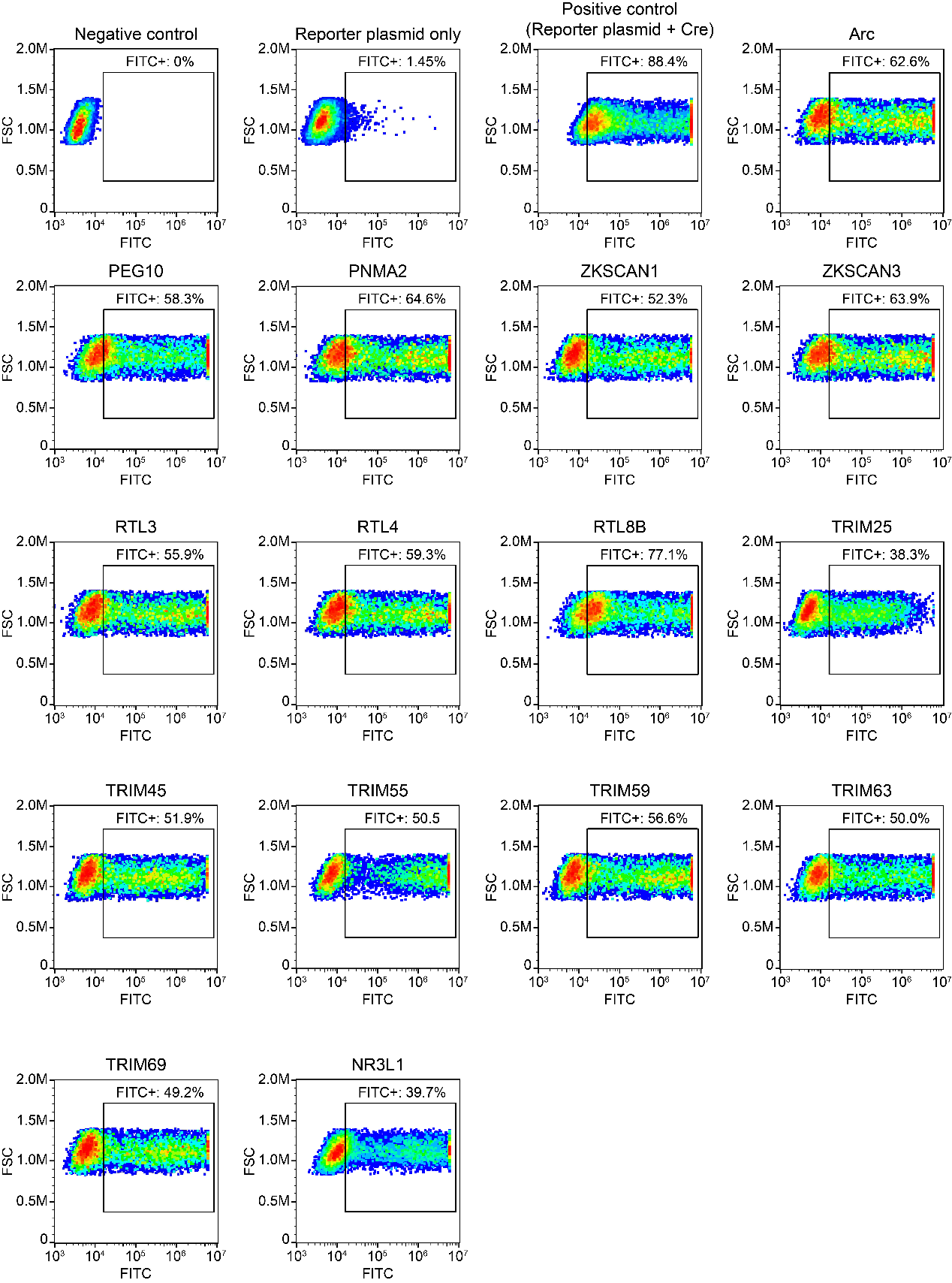
 **Supplementary Fig.15. One batch of flow cytometry results, analyzed in FlowJo v10.9.0.**

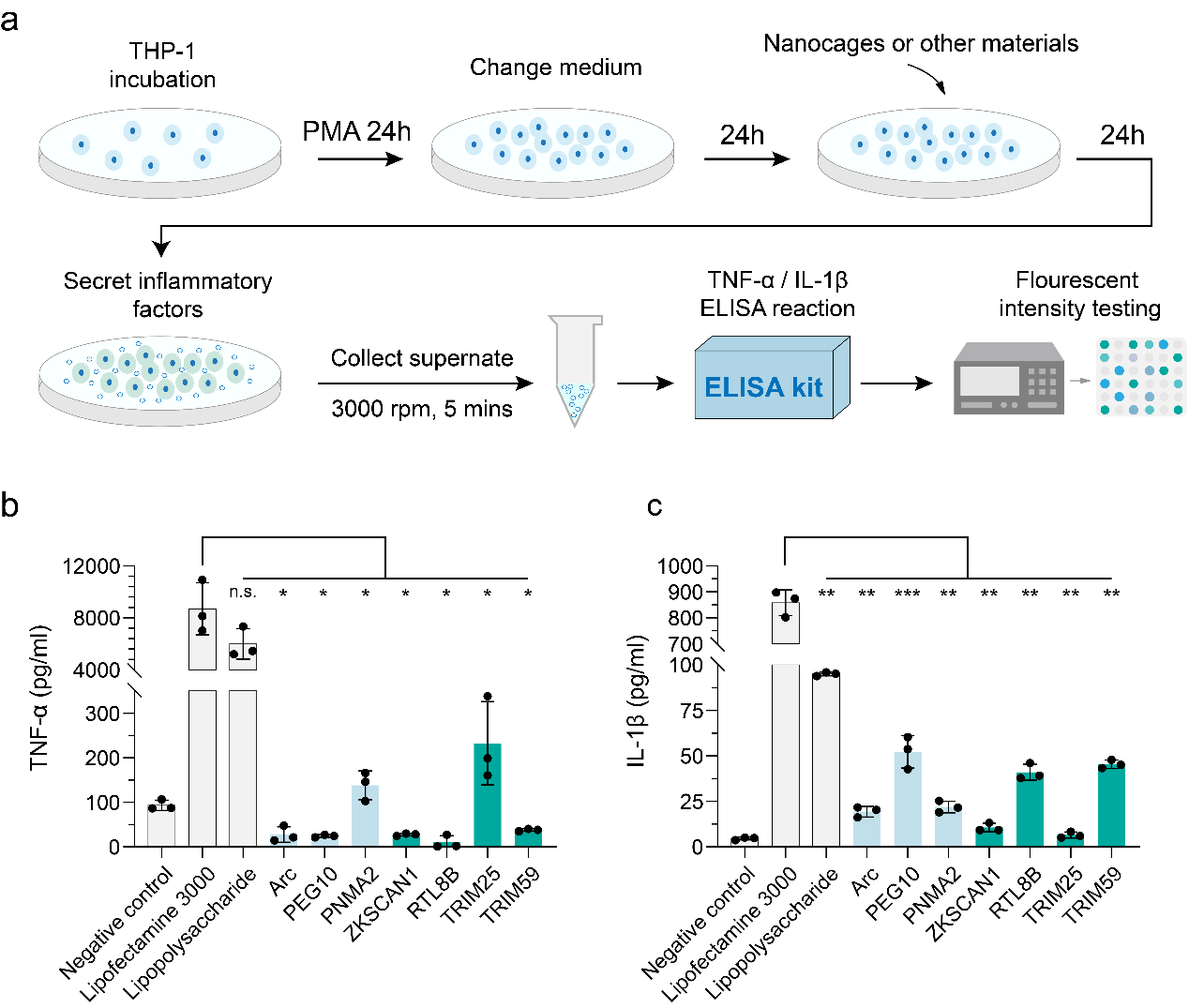
**Supplementary Fig.16. Innate immunogenicity test via THP-1 stimulation. a**, Protocol diagram of innate immunogenicity test (details in Methods). **b**, TNF-α concentration of all groups. **c**, IL-1β concentration of all groups. *n* = 3 independent experiments, mean ± SD. Statistics by unpaired two-tailed Student’s t-test; P≥0.05, not significant, *P < 0.05, **P < 0.01, ***P < 0.001.

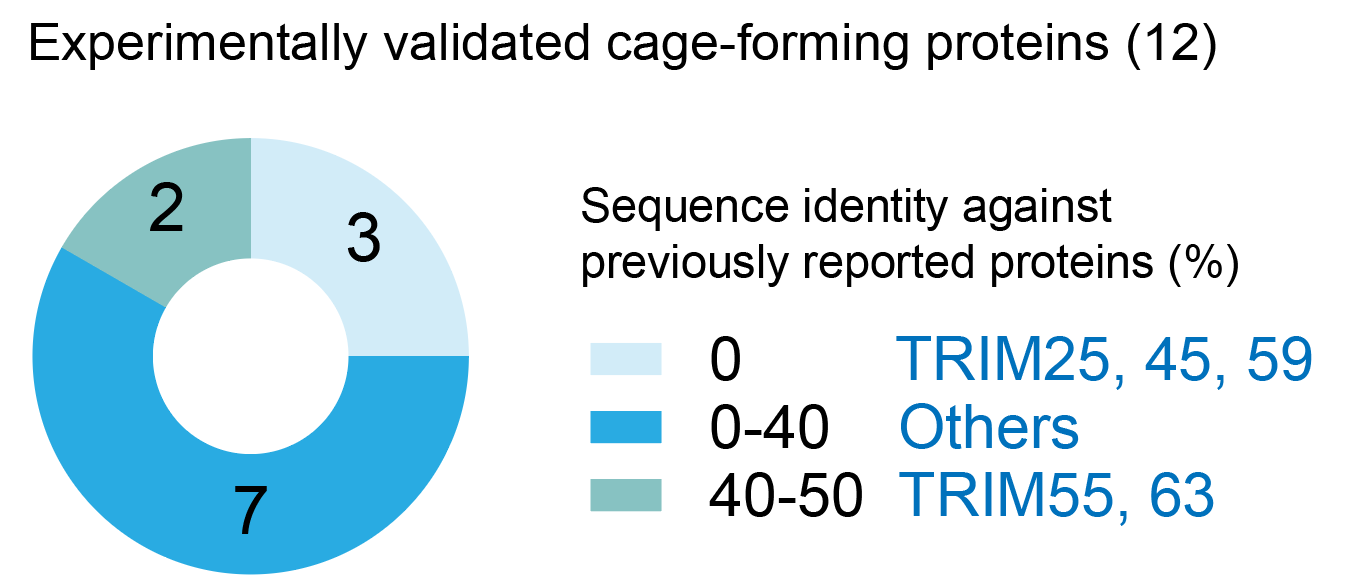

**Supplementary Fig.17. Experimentally validated cage-forming proteins sequence identity against previously reported proteins, tested via Diamond BLASTP.**

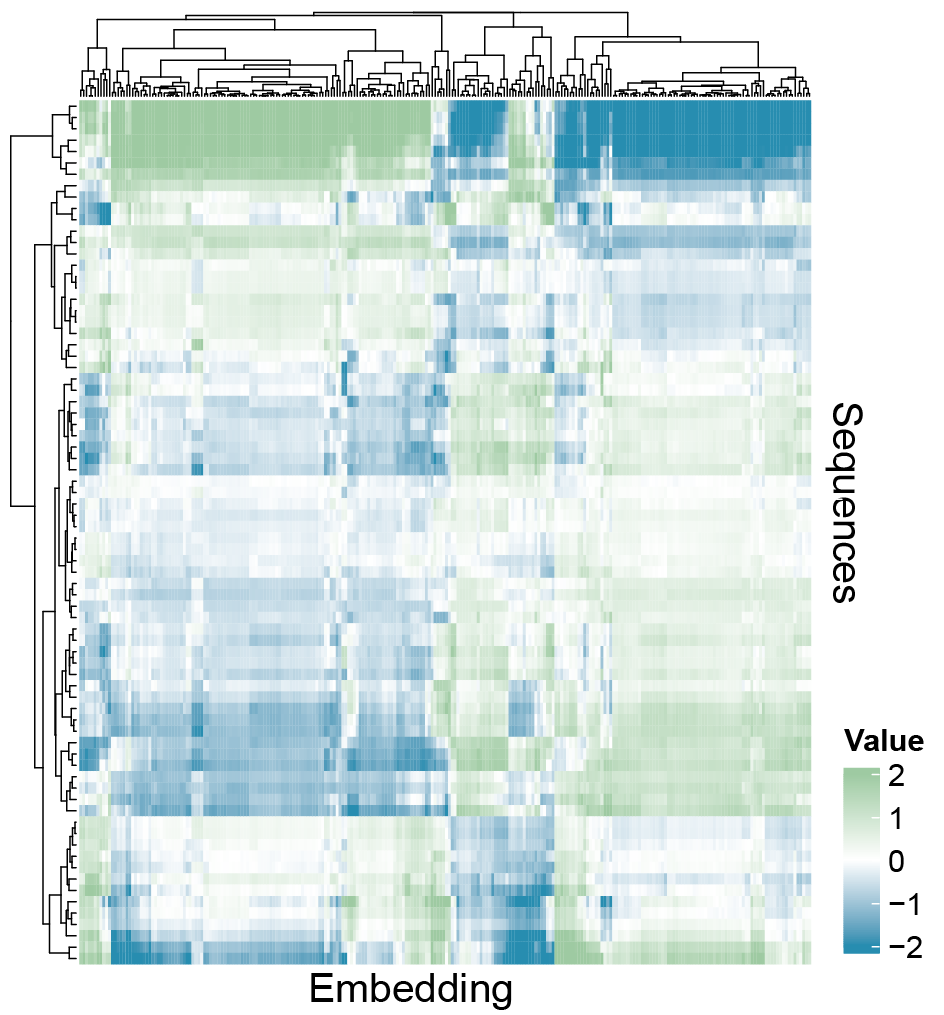

**Supplementary Fig.18. Embedding heatmap of all TRIM family candidates generated by the 1D-CNN model.**

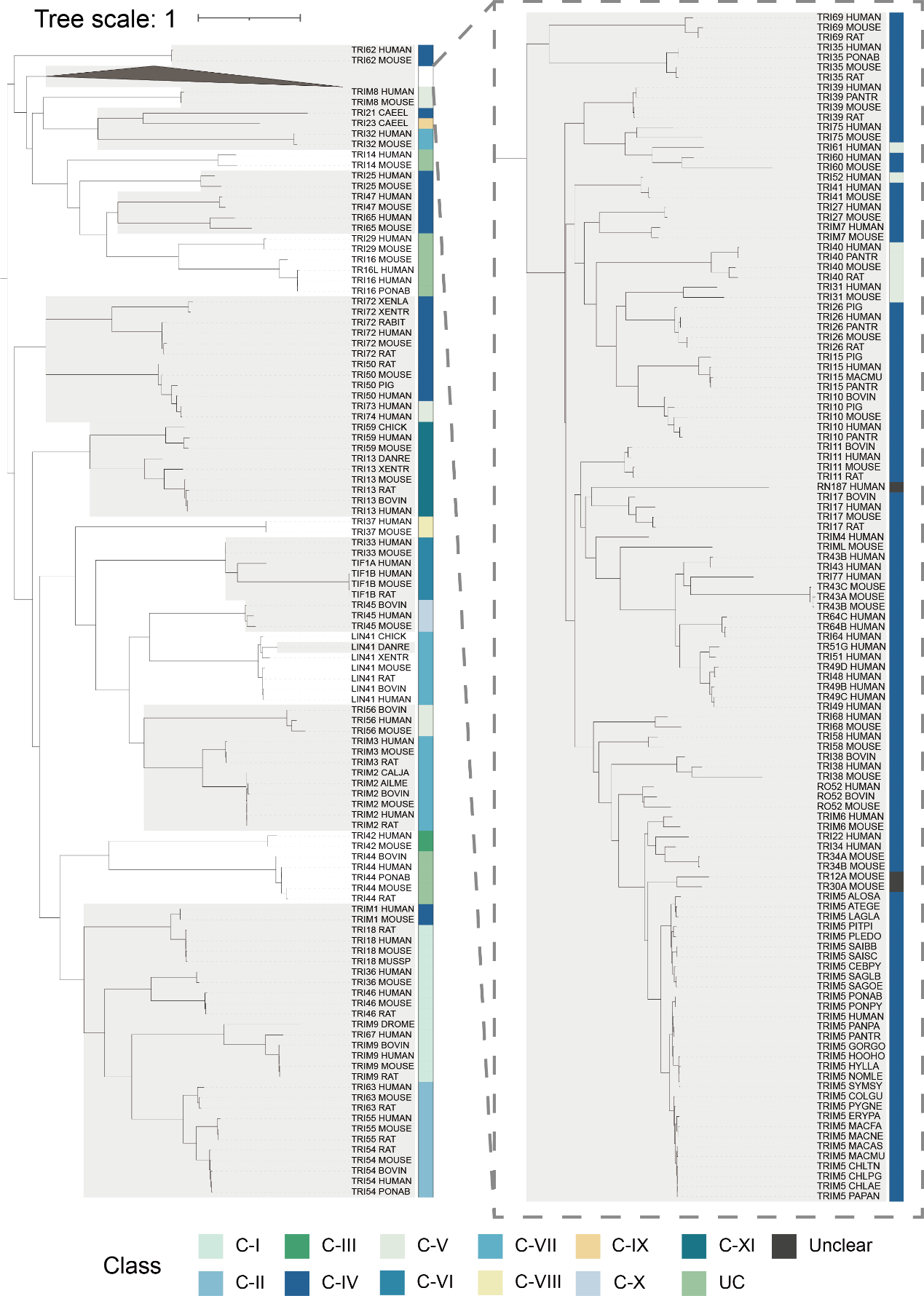

**Supplementary Fig.19.** **Full phylogenetic tree of Figure 5 B.** Distinct clustering patterns correspond to TRIM subfamily classifications, with class labels indicated in Figure 5A.

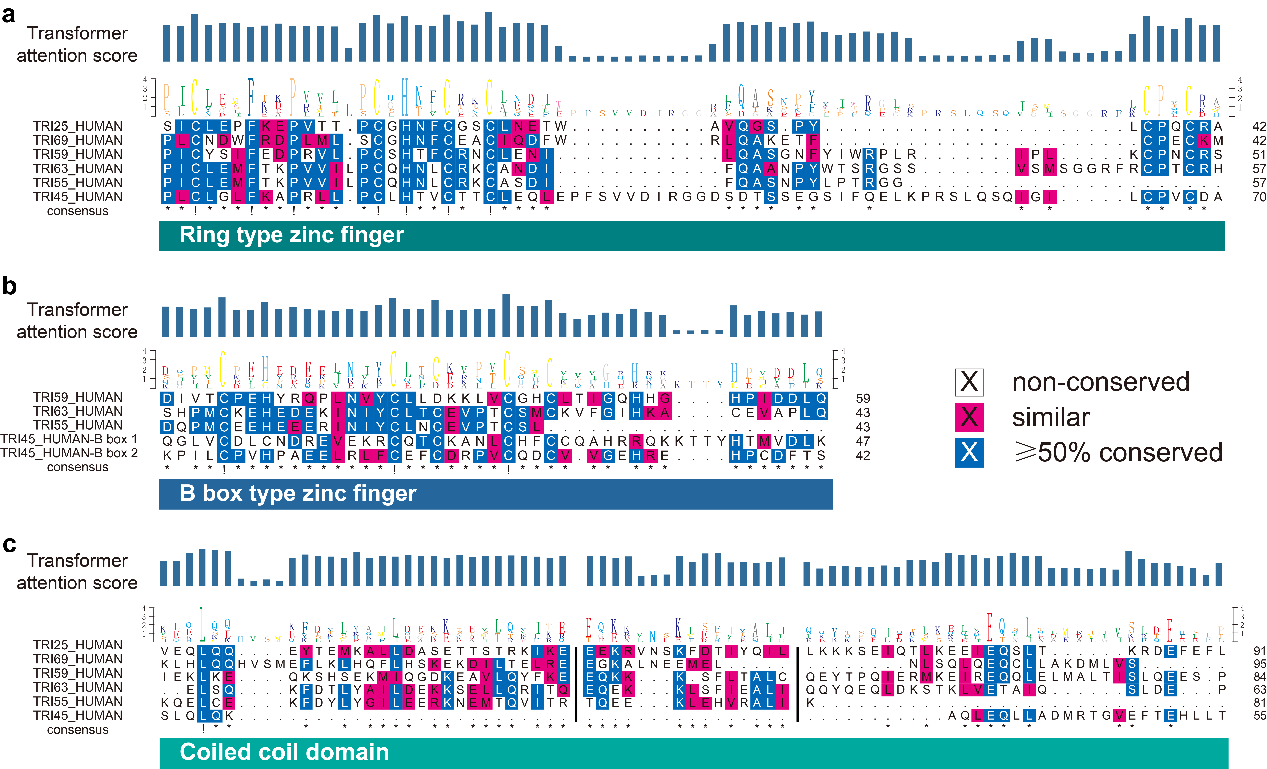

**Supplementary Fig.20.** **Distribution of attention scores of transformer-based deep learning model across the three most frequent domains. a,** ring-type zinc finger. **b**, B-box type zinc finger. **c**, coiled-coil domain.

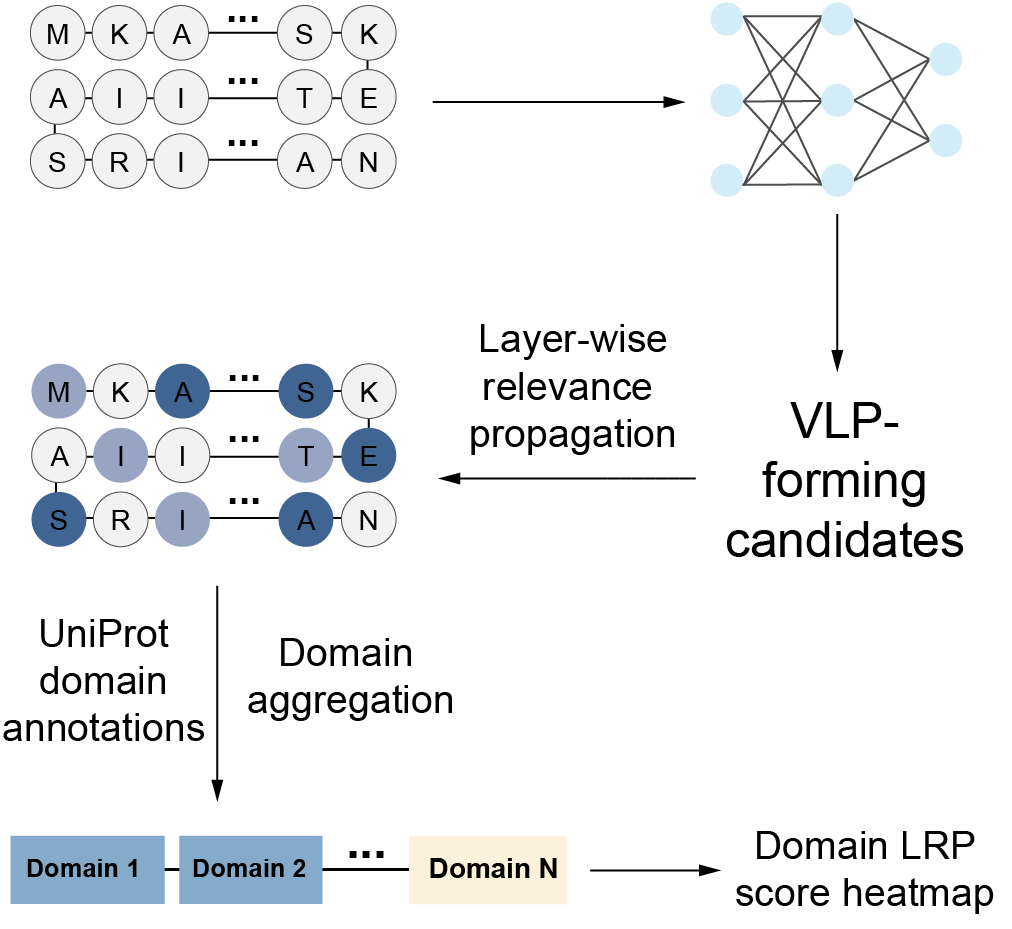

**Supplementary Fig.21.** **Workflow of Layer-wise Relevance Propagation (LRP) analysis.** Protein amino acid sequences were fed into a deep learning model to identify VLP-forming candidates. LRP was used to assess the relevance score of each amino acid. Domain LRP scores were then calculated based on the amino acid LRP scores, incorporating UniProt domain features.

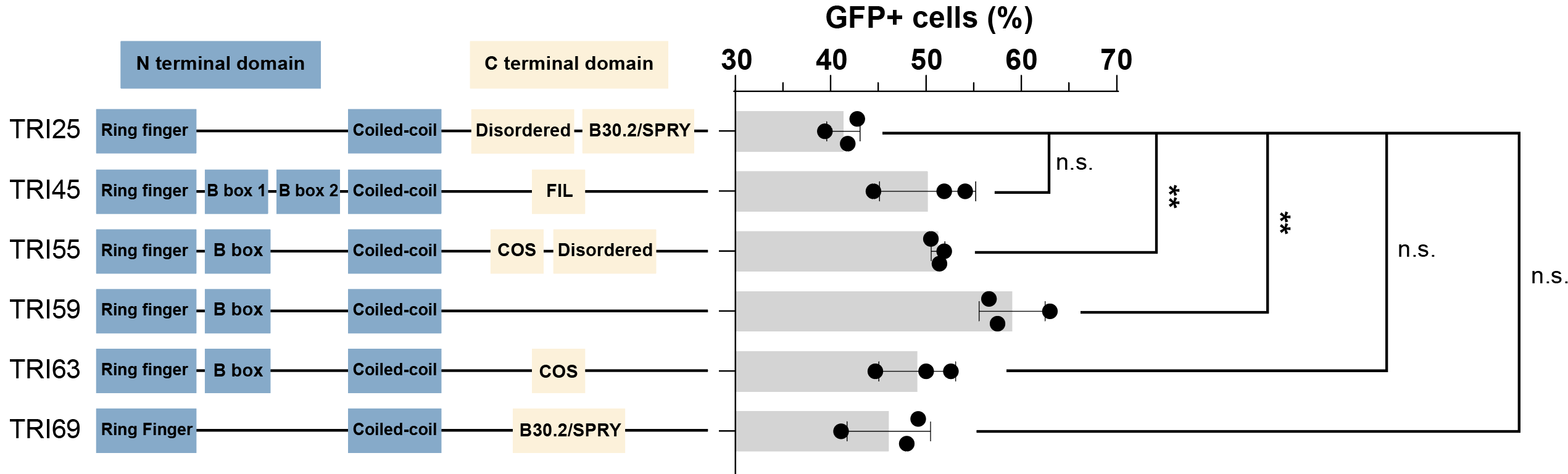

**Supplementary Fig.22.** **Gene structure and transduction efficiency of TRIM proteins.** Left: Gene structure of experiment-validated TRIM family proteins. Right: transduction efficiency, same as shown in Figure 4F. *n* = 3 independent experiments, mean ± SD. Statistics by unpaired two-tailed Student’s t-test; P≥0.05, not significant, *P < 0.05, **P < 0.01, ***P < 0.001.

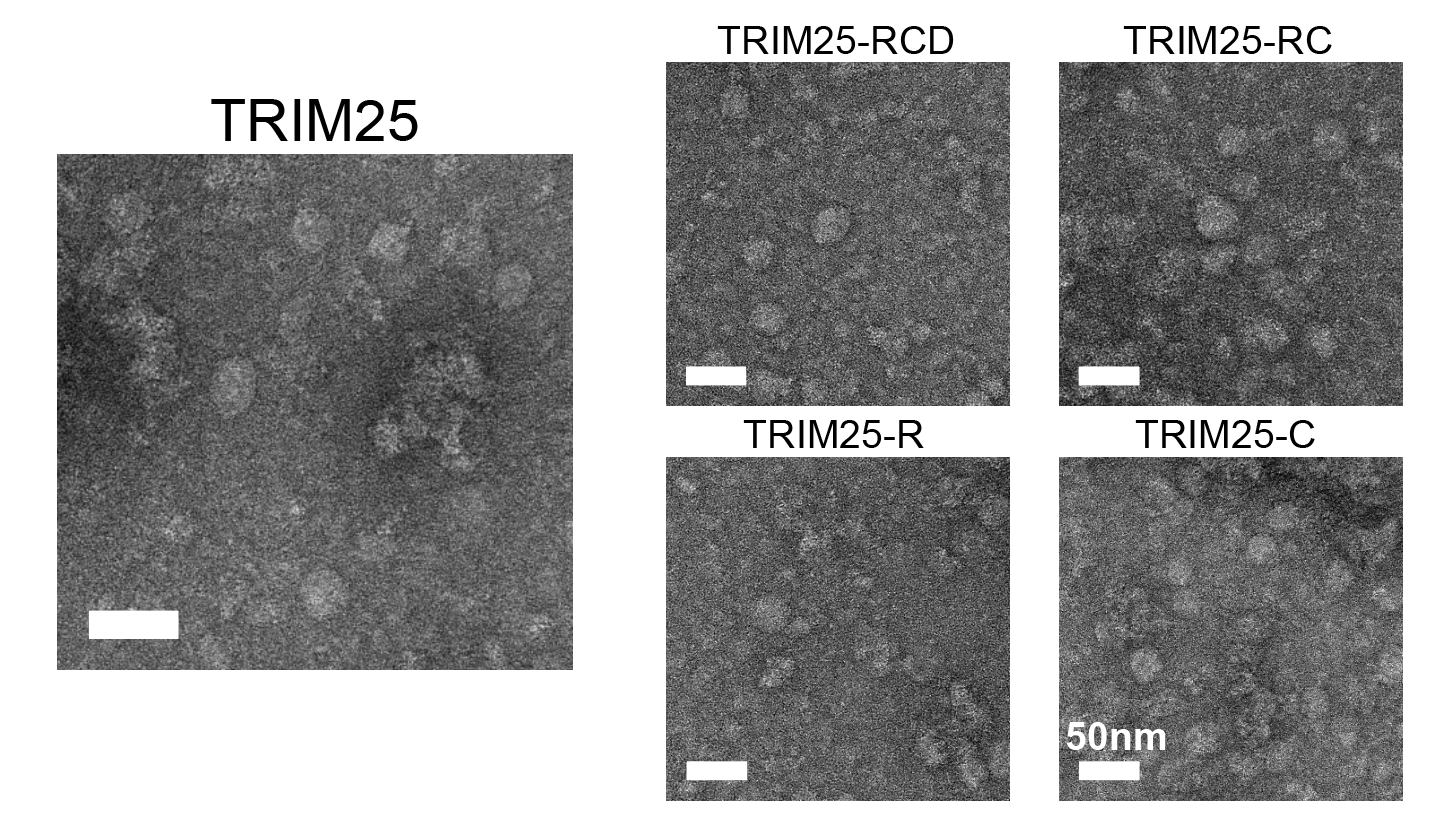

**Supplementary Fig.23.** **Negative-stain TEM imaging of TRIM25 construct-derived nanocages. Representative micrographs showing capsid-like particles.** Scale bar, 50 nm.

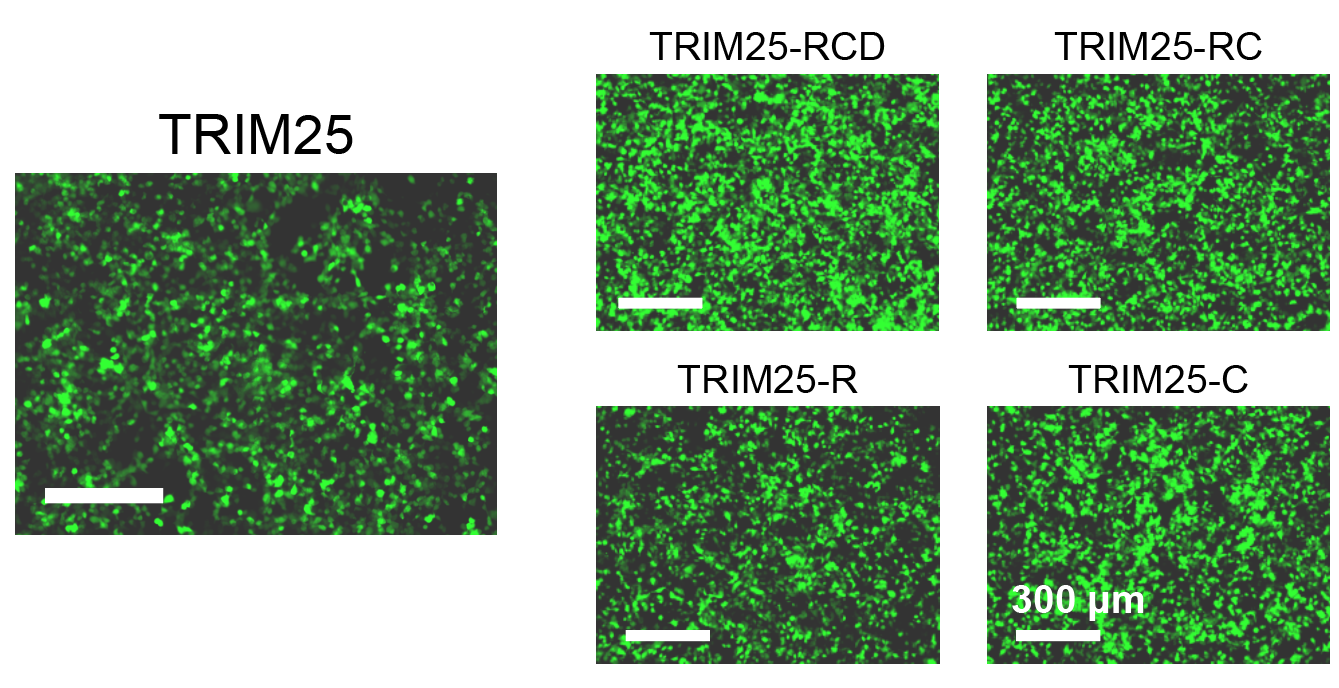

**Supplementary Fig.24.** **Fluorescence microscopy of cells infected with TRIM25 construct-derived nanocages.** Scale bar, 300 μm.

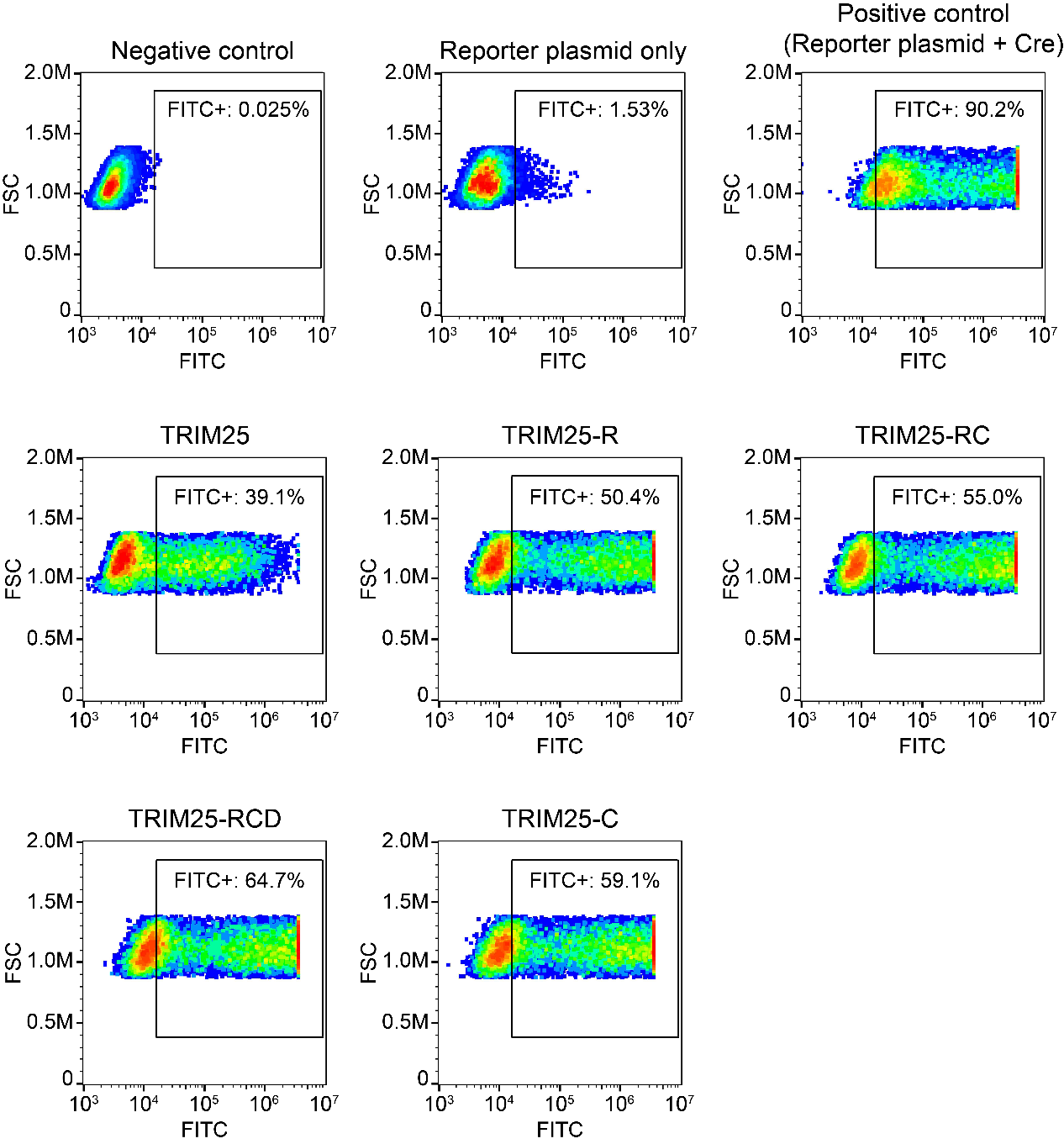

**Supplementary Fig.25.** **One batch of flow cytometry results of TRIM25 construct-derived nanocages, analyzed in FlowJo v10.9.0.**

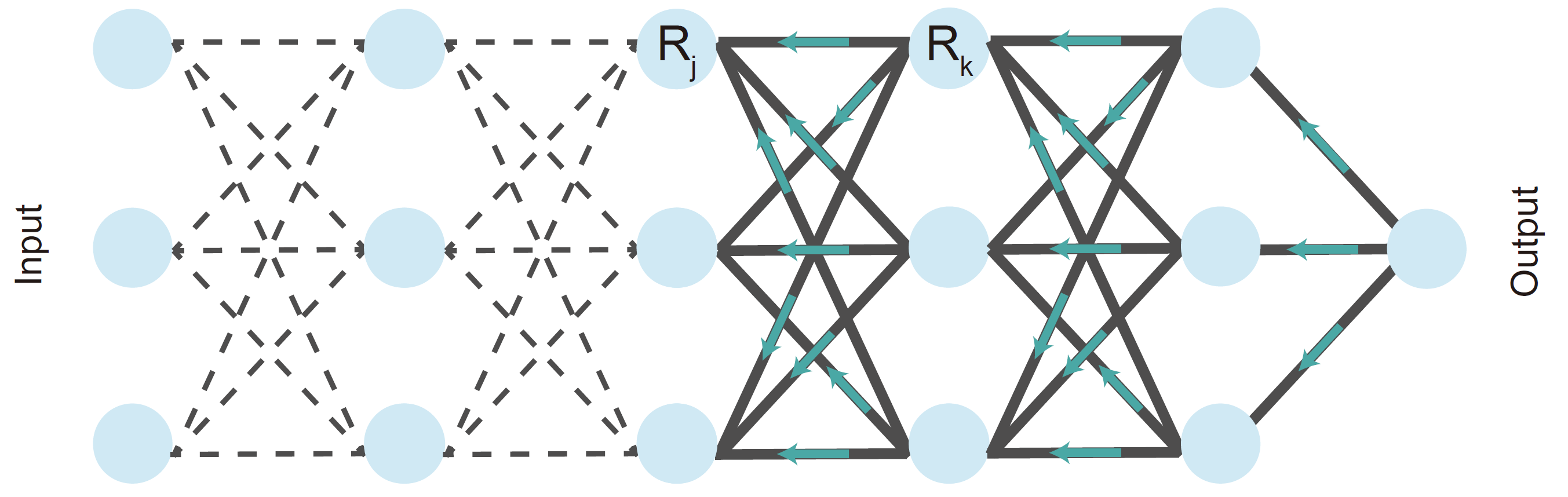

**Supplementary Fig.26.** **Illustration of the LRP procedure.** Each neuron redistributes to the lower layer as much as it has received from the higher layer ^14^.

**Supplemental tables**

**Supplemental Table 1. Candidate protein sequences used in this study.**

|  | Sequences |
| --- | --- |
| ZKSC1_HUMAN | MMTAESREATGLSPQAAQEKDGIVIVKVEEEDEEDHMWGQDSTLQDTPPPDPEIFRQRFRRFCYQNTFGPREALSRLKELCHQWLRPEINTKEQILELLVLEQFLSILPKELQVWLQEYRPDSGEEAVTLLEDLELDLSGQQVPGQVHGPEMLARGMVPLDPVQESSSFDLHHEATQSHFKHSSRKPRLLQSRALPAAHIPAPPHEGSPRDQAMASALFTADSQAMVKIEDMAVSLILEEWGCQNLARRNLSRDNRQENYGSAFPQGGENRNENEESTSKAETSEDSASRGETTGRSQKEFGEKRDQEGKTGERQQKNPEEKTRKEKRDSGPAIGKDKKTITGERGPREKGKGLGRSFSLSSNFTTPEEVPTGTKSHRCDECGKCFTRSSSLIRHKIIHTGEKPYECSECGKAFSLNSNLVLHQRIHTGEKPHECNECGKAFSHSSNLILHQRIHSGEKPYECNECGKAFSQSSDLTKHQRIHTGEKPYECSECGKAFNRNSYLILHRRIHTREKPYKCTKCGKAFTRSSTLTLHHRIHARERASEYSPASLDAFGAFLKSCV |
| ZKSC3_HUMAN | MARELSESTALDAQSTEDQMELLVIKVEEEEAGFPSSPDLGSEGSRERFRGFRYPEAAGPREALSRLRELCRQWLQPEMHSKEQILELLVLEQFLTILPGNLQSWVREQHPESGEEVVVLLEYLERQLDEPAPQVSGVDQGQELLCCKMALLTPAPGSQSSQFQLMKALLKHESVGSQPLQDRVLQVPVLAHGGCCREDKVVASRLTPESQGLLKVEDVALTLTPEWTQQDSSQGNLCRDEKQENHGSLVSLGDEKQTKSRDLPPAEELPEKEHGKISCHLREDIAQIPTCAEAGEQEGRLQRKQKNATGGRRHICHECGKSFAQSSGLSKHRRIHTGEKPYECEECGKAFIGSSALVIHQRVHTGEKPYECEECGKAFSHSSDLIKHQRTHTGEKPYECDDCGKTFSQSCSLLEHHRIHTGEKPYQCSMCGKAFRRSSHLLRHQRIHTGDKNVQEPEQGEAWKSRMESQLENVETPMSYKCNECERSFTQNTGLIEHQKIHTGEKPYQCNACGKGFTRISYLVQHQRSHVGKNILSQ |
| TRI69_HUMAN | MEVSTNPSSNIDPGDYVEMNDSITHLPSKVVIQDITMELHCPLCNDWFRDPLMLSCGHNFCEACIQDFWRLQAKETFCPECKMLCQYNNCTFNPVLDKLVEKIKKLPLLKGHPQCPEHGENLKLFSKPDGKLICFQCKDARLSVGQSKEFLQISDAVHFFTEELAIQQGQLETTLKELQTLRNMQKEAIAAHKENKLHLQQHVSMEFLKLHQFLHSKEKDILTELREEGKALNEEMELNLSQLQEQCLLAKDMLVSIQAKTEQQNSFDFLKDITTLLHSLEQGMKVLATRELISRKLNLGQYKGPIQYMVWREMQDTLCPGLSPLTLDPKTAHPNLVLSKSQTSVWHGDIKKIMPDDPERFDSSVAVLGSRGFTSGKWYWEVEVAKKTKWTVGVVRESIIRKGSCPLTPEQGFWLLRLRNQTDLKALDLPSFSLTLTNNLDKVGIYLDYEGGQLSFYNAKTMTHIYTFSNTFMEKLYPYFCPCLNDGGENKEPLHILHPQ |
| TRI63_HUMAN | MDYKSSLIQDGNPMENLEKQLICPICLEMFTKPVVILPCQHNLCRKCANDIFQAANPYWTSRGSSVSMSGGRFRCPTCRHEVIMDRHGVYGLQRNLLVENIIDIYKQECSSRPLQKGSHPMCKEHEDEKINIYCLTCEVPTCSMCKVFGIHKACEVAPLQSVFQGQKTELNNCISMLVAGNDRVQTIITQLEDSRRVTKENSHQVKEELSQKFDTLYAILDEKKSELLQRITQEQEKKLSFIEALIQQYQEQLDKSTKLVETAIQSLDEPGGATFLLTAKQLIKSIVEASKGCQLGKTEQGFENMDFFTLDLEHIADALRAIDFGTDEEEEEFIEEEDQEEEESTEGKEEGHQ |
| TRI59_HUMAN | MHNFEEELTCPICYSIFEDPRVLPCSHTFCRNCLENILQASGNFYIWRPLRIPLKCPNCRSITEIAPTGIESLPVNFALRAIIEKYQQEDHPDIVTCPEHYRQPLNVYCLLDKKLVCGHCLTIGQHHGHPIDDLQSAYLKEKDTPQKLLEQLTDTHWTDLTHLIEKLKEQKSHSEKMIQGDKEAVLQYFKELNDTLEQKKKSFLTALCDVGNLINQEYTPQIERMKEIREQQLELMALTISLQEESPLKFLEKVDDVRQHVQILKQRPLPEVQPVEIYPRVSKILKEEWSRTEIGQIKNVLIPKMKISPKRMSCSWPGKDEKEVEFLKILNIVVVTLISVILMSILFFNQHIITFLSEITLIWFSEASLSVYQSLSNSLHKVKNILCHIFYLLKEFVWKIVSH |
| TRI55_HUMAN | MSASLNYKSFSKEQQTMDNLEKQLICPICLEMFTKPVVILPCQHNLCRKCASDIFQASNPYLPTRGGTTMASGGRFRCPSCRHEVVLDRHGVYGLQRNLLVENIIDIYKQESTRPEKKSDQPMCEEHEEERINIYCLNCEVPTCSLCKVFGAHKDCQVAPLTHVFQRQKSELSDGIAILVGSNDRVQGVISQLEDTCKTIEECCRKQKQELCEKFDYLYGILEERKNEMTQVITRTQEEKLEHVRALIKKYSDHLENVSKLVESGIQFMDEPEMAVFLQNAKTLLKKISEASKAFQMEKIEHGYENMNHFTVNLNREEKIIREIDFYREDEDEEEEEGGEGEKEGEGEVGGEAVEVEEVENVQTEFPGEDENPEKASELSQVELQAAPGALPVSSPEPPPALPPAADAPVTQGEVVPTGSEQTTESETPVPAAAETADPLFYPSWYKGQTRKATTNPPCTPGSEGLGQIGPPGSEDSNVRKAEVAAAAASERAAVSGKETSAPAATSQIGFEAPPLQGQAAAPASGSGADSEPARHIFSFSWLNSLNE |
| TRI45_HUMAN | MSENRKPLLGFVSKLTSGTALGNSGKTHCPLCLGLFKAPRLLPCLHTVCTTCLEQLEPFSVVDIRGGDSDTSSEGSIFQELKPRSLQSQIGILCPVCDAQVDLPMGGVKALTIDHLAVNDVMLESLRGEGQGLVCDLCNDREVEKRCQTCKANLCHFCCQAHRRQKKTTYHTMVDLKDLKGYSRIGKPILCPVHPAEELRLFCEFCDRPVCQDCVVGEHREHPCDFTSNVIHKHGDSVWELLKGTQPHVEALEEALAQIHIINSALQKRVEAVAADVRTFSEGYIKAIEEHRDKLLKQLEDIRAQKENSLQLQKAQLEQLLADMRTGVEFTEHLLTSGSDLEILITKRVVVERLRKLNKVQYSTRPGVNDKIRFCPQEKAGQCRGYEIYGTINTKEVDPAKCVLQGEDLHRAREKQTASFTLLCKDAAGEIMGRGGDNVQVAVVPKDKKDSPVRTMVQDNKDGTYYISYTPKEPGVYTVWVCIKEQHVQGSPFTVMVRRKHRPHSGVFHCCTFCSSGGQKTARCACGGTMPGGYLGCGHGHKGHPGHPHWSCCGKFNEKSECTWTGGQSAPRSLLRTVAL |
| TRI25_HUMAN | MAELCPLAEELSCSICLEPFKEPVTTPCGHNFCGSCLNETWAVQGSPYLCPQCRAVYQARPQLHKNTVLCNVVEQFLQADLAREPPADVWTPPARASAPSPNAQVACDHCLKEAAVKTCLVCMASFCQEHLQPHFDSPAFQDHPLQPPVRDLLRRKCSQHNRLREFFCPEHSECICHICLVEHKTCSPASLSQASADLEATLRHKLTVMYSQINGASRALDDVRNRQQDVRMTANRKVEQLQQEYTEMKALLDASETTSTRKIKEEEKRVNSKFDTIYQILLKKKSEIQTLKEEIEQSLTKRDEFEFLEKASKLRGISTKPVYIPEVELNHKLIKGIHQSTIDLKNELKQCIGRLQEPTPSSGDPGEHDPASTHKSTRPVKKVSKEEKKSKKPPPVPALPSKLPTFGAPEQLVDLKQAGLEAAAKATSSHPNSTSLKAKVLETFLAKSRPELLEYYIKVILDYNTAHNKVALSECYTVASVAEMPQNYRPHPQRFTYCSQVLGLHCYKKGIHYWEVELQKNNFCGVGICYGSMNRQGPESRLGRNSASWCVEWFNTKISAWHNNVEKTLPSTKATRVGVLLNCDHGFVIFFAVADKVHLMYKFRVDFTEALYPAFWVFSAGATLSICSPK |
| RTL8C_HUMAN | MDGRVQLIKALLALPIRPATRRWRNPIPFPETFDGDTDRLPEFIVQTGSYMFVDENTFSSDALKVTFLITRLTGPALQWVIPYIKKESPLLNDYRGFLAEMKRVFGWEEDEDF |
| RTL8B_HUMAN | MEGRVQLMKALLARPLRPAARRWRNPIPFPETFDGDTDRLPEFIVQTSSYMFVDENTFSNDALKVTFLITRLTGPALQWVIPYIKKESPLLSDYRGFLAEMKRVFGWEEDEDF |
| RTL4_HUMAN | MEKCTKSSSTMQVEPSFLQAENLILRLQMQHPTTENTAKRGQVMPALATTVMPVPYSLEHLTQFHGDPANCSEFLTQVTTYLTALQISNPANDAQIKLFFDYLSQQLESCGIISGPDKSTLLKQYENLILEFQQSFGKPTKQEINPLMNAKFDKGDNSSQQDPATFHLLAQNLICNETNQSGQFEKALADPNQDEESVTDMMDNLPDLITQCIQLDKKHSDRPELLQSETQLPLLASLIQHQALFSPTDPPPKKGPIQLREGQLPLTPAKRARQQETQLCLYCSQSGHFTRDCLAKRSRAPATTNNTAHQ |
| RTL3_HUMAN | MVEDLAASYIVLKLENEIRQAQVQWLMEENAALQAQIPELQKSQAAKEYDLLRKSSEAKEPQKLPEHMNPPAAWEAQKTPEFKEPQKPPEPQDLLPWEPPAAWELQEAPAAPESLAPPATRESQKPPMAHEIPTVLEGQGPANTQDATIAQEPKNSEPQDPPNIEKPQEAPEYQETAAQLEFLELPPPQEPLEPSNAQEFLELSAAQESLEGLIVVETSAASEFPQAPIGLEATDFPLQYTLTFSGDSQKLPEFLVQLYSYMRVRGHLYPTEAALVSFVGNCFSGRAGWWFQLLLDIQSPLLEQCESFIPVLQDTFDNPENMKDANQCIHQLCQGEGHVATHFHLIAQELNWDESTLWIQFQEGLASSIQDELSHTSPATNLSDLITQCISLEEKPDPNPLGKSSSAEGDGPESPPAENQPMQAAINCPHISEAEWVRWHKGRLCLYCGYPGHFARDCPVKPHQALQAGNIQACQ |
| RN186_HUMAN | MACTKTLQQSQPISAGATTTTTAVAPAGGHSGSTECDLECLVCREPYSCPRLPKLLACQHAFCAICLKLLLCVQDNTWSITCPLCRKVTAVPGGLICSLRDHEAVVGQLAQPCTEVSLCPQGLVDPADLAAGHPSLVGEDGQDEVSANHVAARRLAAHLLLLALLIILIGPFIYPGVLRWVLTFIIALALLMSTLFCCLPSTRGSCWPSSRTLFCREQKHSHISSIA |
| PNMA2_HUMAN | MALALLEDWCRIMSVDEQKSLMVTGIPADFEEAEIQEVLQETLKSLGRYRLLGKIFRKQENANAVLLELLEDTDVSAIPSEVQGKGGVWKVIFKTPNQDTEFLERLNLFLEKEGQTVSGMFRALGQEGVSPATVPCISPELLAHLLGQAMAHAPQPLLPMRYRKLRVFSGSAVPAPEEESFEVWLEQATEIVKEWPVTEAEKKRWLAESLRGPALDLMHIVQADNPSISVEECLEAFKQVFGSLESRRTAQVRYLKTYQEEGEKVSAYVLRLETLLRRAVEKRAIPRRIADQVRLEQVMAGATLNQMLWCRLRELKDQGPPPSFLELMKVIREEEEEEASFENESIEEPEERDGYGRWNHEGDD |
| PGBD1_HUMAN | MYEALPGPAPENEDGLVKVKEEDPTWEQVCNSQEGSSHTQEICRLRFRHFCYQEAHGPQEALAQLRELCHQWLRPEMHTKEQIMELLVLEQFLTILPKELQPCVKTYPLESGEEAVTVLENLETGSGDTGQQASVYIQGQDMHPMVAEYQGVSLECQSLQLLPGITTLKCEPPQRPQGNPQEVSGPVPHGSAHLQEKNPRDKAVVPVFNPVRSQTLVKTEEETAQAVAAEKWSHLSLTRRNLCGNSAQETVMSLSPMTEEIVTKDRLFKAKQETSEEMEQSGEASGKPNRECAPQIPCSTPIATERTVAHLNTLKDRHPGDLWARMHISSLEYAAGDITRKGRKKDKARVSELLQGLSFSGDSDVEKDNEPEIQPAQKKLKVSCFPEKSWTKRDIKPNFPSWSALDSGLLNLKSEKLNPVELFELFFDDETFNLIVNETNNYASQKNVSLEVTVQEMRCVFGVLLLSGFMRHPRREMYWEVSDTDQNLVRDAIRRDRFELIFSNLHFADNGHLDQKDKFTKLRPLIKQMNKNFLLYAPLEEYYCFDKSMCECFDSDQFLNGKPIRIGYKIWCGTTTQGYLVWFEPYQEESTMKVDEDPDLGLGGNLVMNFADVLLERGQYPYHLCFDSFFTSVKLLSALKKKGVRATGTIRENRTEKCPLMNVEHMKKMKRGYFDFRIEENNEIILCRWYGDGIISLCSNAVGIEPVNEVSCCDADNEEIPQISQPSIVKVYDECKEGVAKMDQIISKYRVRIRSKKWYSILVSYMIDVAMNNAWQLHRACNPGASLDPLDFRRFVAHFYLEHNAHLSD |
| NR3L1_HUMAN | MTWRAAASTCAALLILLWALTTEGDLKVEMMAGGTQITPLNDNVTIFCNIFYSQPLNITSMGITWFWKSLTFDKEVKVFEFFGDHQEAFRPGAIVSPWRLKSGDASLRLPGIQLEEAGEYRCEVVVTPLKAQGTVQLEVVASPASRLLLDQVGMKENEDKYMCESSGFYPEAINITWEKQTQKFPHPIEISEDVITGPTIKNMDGTFNVTSCLKLNSSQEDPGTVYQCVVRHASLHTPLRSNFTLTAARHSLSETEKTDNFSIHWWPISFIGVGLVLLIVLIPWKKICNKSSSAYTPLKCILKHWNSFDTQTLKKEHLIFFCTRAWPSYQLQDGEAWPPEGSVNINTIQQLDVFCRQEGKWSEVPYVQAFFALRDNPDLCQCCRIDPALLTVTSGKSIDDNSTKSEKQTPREHSDAVPDAPILPVSPIWEPPPATTSTTPVLSSQPPTLLLPLQ |
| ARC_HUMAN | MELDHRTSGGLHAYPGPRGGQVAKPNVILQIGKCRAEMLEHVRRTHRHLLAEVSKQVERELKGLHRSVGKLESNLDGYVPTSDSQRWKKSIKACLCRCQETIANLERWVKREMHVWREVFYRLERWADRLESTGGKYPVGSESARHTVSVGVGGPESYCHEADGYDYTVSPYAITPPPAAGELPGQEPAEAQQYQPWVPGEDGQPSPGVDTQIFEDPREFLSHLEEYLRQVGGSEEYWLSQIQNHMNGPAKKWWEFKQGSVKNWVEFKKEFLQYSEGTLSREAIQRELDLPQKQGEPLDQFLWRKRDLYQTLYVDADEEEIIQYVVGTLQPKLKRFLRHPLPKTLEQLIQRGMEVQDDLEQAAEPAGPHLPVEDEAETLTPAPNSESVASDRTQPE |
| PEG10_HUMAN | MTERRRDELSEEINNLREKVMKQSEENNNLQSQVQKLTEENTTLREQVEPTPEDEDDDIELRGAAAAAAPPPPIEEECPEDLPEKFDGNPDMLAPFMAQCQIFMEKSTRDFSVDRVRVCFVTSMMTGRAARWASAKLERSHYLMHNYPAFMMEMKHVFEDPQRREVAKRKIRRLRQGMGSVIDYSNAFQMIAQDLDWNEPALIDQYHEGLSDHIQEELSHLEVAKSLSALIGQCIHIERRLARAAAARKPRSPPRALVLPHIASHHQVDPTEPVGGARMRLTQEEKERRRKLNLCLYCGTGGHYADNCPAKASKSSPAGKLPGPAVEGPSATGPEIIRSPQDDASSPHLQVMLQIHLPGRHTLFVRAMIDSGASGNFIDHEYVAQNGIPLRIKDWPILVEAIDGRPIASGPVVHETHDLIVDLGDHREVLSFDVTQSPFFPVVLGVRWLSTHDPNITWSTRSIVFDSEYCRYHCRMYSPIPPSLPPPAPQPPLYYPVDGYRVYQPVRYYYVQNVYTPVDEHVYPDHRLVDPHIEMIPGAHSIPSGHVYSLSEPEMAALRDFVARNVKDGLITPTIAPNGAQVLQVKRGWKLQVSYDCRAPNNFTIQNQYPRLSIPNLEDQAHLATYTEFVPQIPGYQTYPTYAAYPTYPVGFAWYPVGRDGQGRSLYVPVMITWNPHWYRQPPVPQYPPPQPPPPPPPPPPPPSYSTL |

**Supplemental Table 2. Summary of experimentally validated nanocage candidates and their experimental outcomes.**

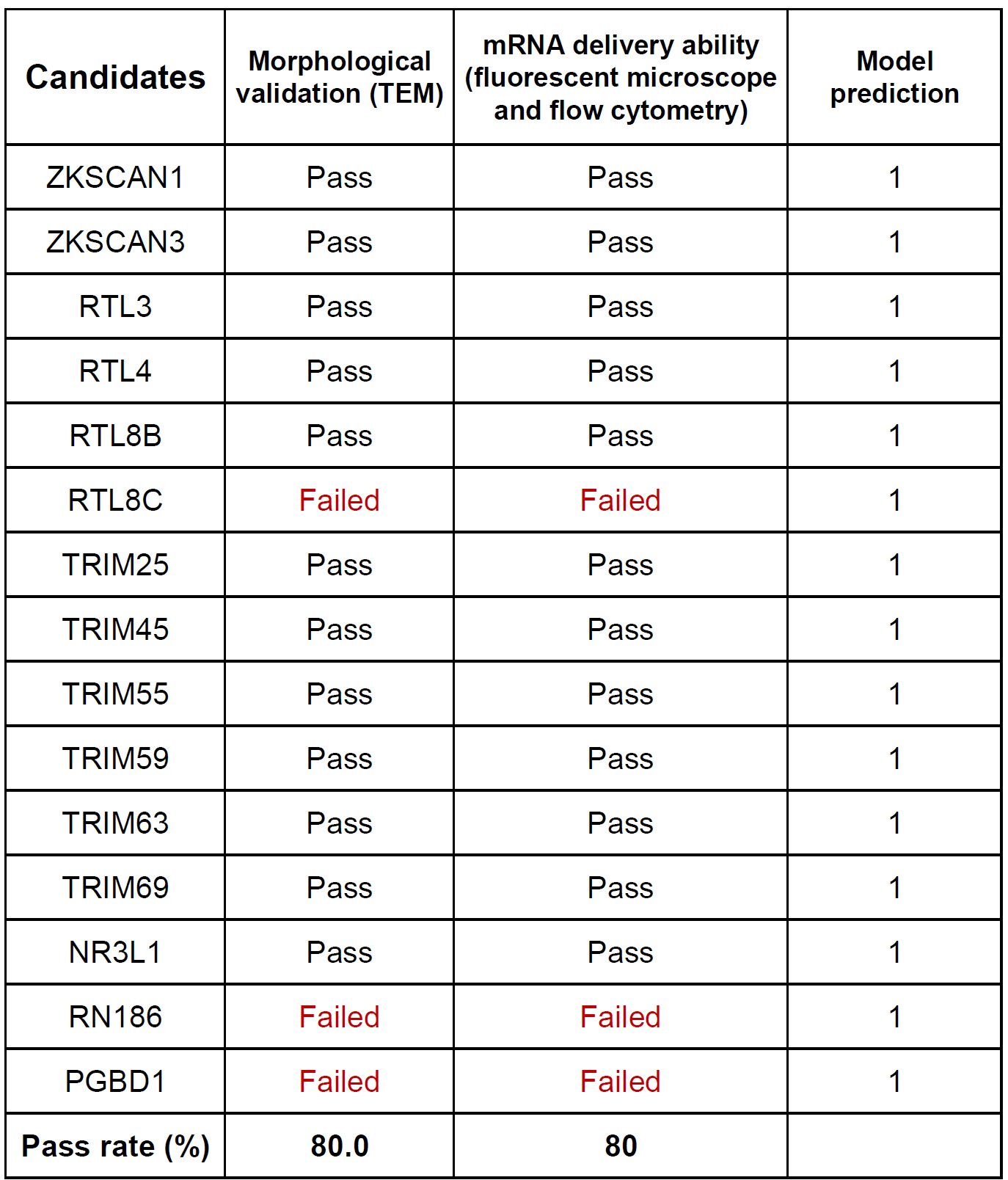

**Supplemental Table 3. Domain label of LRP analysis from UniProt annotations.**

| **Type** | **Protein Domain** | **Label used in this study** |
| --- | --- | --- |
| Coiled coil | Coiled coil | Coiled coil |
| Domain [FT] | COS | COS |
| Domain [FT] | Fibronectin type-III | Fibronectin type-III |
| Domain [FT] | Fibronectin type-III 1 | Fibronectin type-III |
| Domain [FT] | Fibronectin type-III 2 | Fibronectin type-III |
| Domain [FT] | B30.2/SPRY | B30.2/SPRY |
| Domain [FT] | Bromo | Bromo |
| Domain [FT] | Pyrin | Pyrin |
| Domain [FT] | MATH | MATH |
| Repeat | Filamin | Filamin |
| Repeat | NHL | NHL |
| Motif | Nuclear localization signal | Nuclear localization signal |
| Motif | PxVxL motif | PxVxL motif |
| Zinc finger | RING-type; degenerate | RING-type |
| Zinc finger | RING-type | RING-type |
| Zinc finger | RING-type 1; degenerate | RING-type |
| Zinc finger | RING-type 1 | RING-type |
| Zinc finger | RING-type 2; degenerate | RING-type |
| Zinc finger | RING-type 2 | RING-type |
| Zinc finger | B box-type | B box-type |
| Zinc finger | B box-type; atypical | B box-type |
| Zinc finger | B box-type 1 | B box-type |
| Zinc finger | B box-type 1; atypical | B box-type |
| Zinc finger | B box-type 1; degenerate | B box-type |
| Zinc finger | B box-type 2 | B box-type |
| Zinc finger | PHD-type | PHD-type |
| Region | Disordered | Disordered |
| Region | Leucine zipper alpha helical coiled-coil region | Coiled coil |
| Region | Required for proximal axon localization, axon formation and migration | Related to cytoskeleton |
| Region | Required for microtubule association, proximal axon localization and axon formation | Related to cytoskeleton |
| Region | Interaction with MAGEC2 | Anti cancer |
| Region | Involved in binding PPP1CA | Anti viral |
| Region | Nuclear receptor binding site (NRBS) | Related to nuclear |
| Region | Interaction with histone H3 that is not methylated at 'Lys-4' (H3K4me0) | Related to histone |
| Region | Interaction with histone H3 that is acetylated at 'Lys-23' (H3K23ac) | Related to histone |
| Region | Interaction with RELA | Anti cancer |
| Region | Interaction with KIF21B | Related to cytoskeleton |
| Region | Interaction with PER2 | Anti cancer |
| Region | ARF-like | Related to cytoskeleton |
| Region | Required for interaction with GABARAP and for autophagy | Autophagy |
| Region | HP1 box | Related to histone |
| Region | Interaction with influenza A virus NS1 | Anti viral |
| Region | Interaction with CDKN1A | Anti cancer |
| Region | Interaction with TTN | Related to cytoskeleton |
| Region | Mediates microtubule-binding and homooligomerization | Related to cytoskeleton |
| Region | Necessary for nuclear localization | Related to nuclear |
| Region | Important for rapid proteolytic degradation by the proteasome | Protein degradation |
| Region | Necessary for E3 ubiquitin-protein ligase activity and repression of SMAD4 signaling and transcriptional repression | Protein degradation |
| Region | Sumo interaction motif (SIM) | Protein degradation |
| Region | Necessary for oligomerization | Oligomerization |
| Region | Highly hydrophilic | Hydrophilic domain |
| Region | RBCC domain | RBCC domain |
| Region | Required for homotrimerization and induction of pyroptosomes | Homotrimerization |
| Region | Required for RYR2 clustering | RYR2 clustering |
| Region | Amphipathic helix H1 | Amphipathic domain |
| Region | Amphipathic helix H2 | Amphipathic domain |
| Region | Amphipathic helix H3 | Amphipathic domain |
| Region | B-box coiled-coil; BBC | BCC domain |
